# Biosynthetic engineering of the antifungal, anti-MRSA auroramycin

**DOI:** 10.1101/829812

**Authors:** Wan Lin Yeo, Elena Heng, Lee Ling Tan, Yi Wee Lim, Kuan Chieh Ching, De-Juin Tsai, Yi Wun Jhang, Tsai-Ling Lauderdale, Kak-Shan Shia, Huimin Zhao, Ee Lui Ang, Mingzi M. Zhang, Yee Hwee Lim, Fong T. Wong

**Affiliations:** Metabolic Engineering, Functional Molecules & Polymer, Institute of Chemical and Engineering Sciences, A*STAR, Singapore; Molecular Engineering Laboratory, Institute of Bioengineering and Nanotechnology, A*STAR, Singapore; Integrated Bio & Organic Chemistry, Functional Molecules & Polymer, Institute of Chemical and Engineering Sciences, A*STAR, Singapore; National Institute of Infectious Diseases and Vaccinology (DJT & TLL) and Institute of Biotechnology and Pharmaceutical Research (KSS), National Health Research Institutes (NHRI), Taiwan, R.O.C.; Department of Chemical and Biomolecular engineering, Department of Chemistry, Department of Biochemistry, Institute of Molecular and Genomic Medicine, United States; Institute of Molecular and Genomic Medicine, National Health Research Institutes, Taiwan, R.O.C.

**Keywords:** combinatorial biosynthesis, *Streptomyces*, polyene macrolactam

## Abstract

Using an established CRISPR-Cas mediated genome editing technique for streptomycetes, we explored the combinatorial biosynthesis potential of the auroramycin biosynthetic gene cluster in *Streptomyces roseoporous.* Auroramycin is a potent anti-MRSA polyene macrolactam. In addition, it also displays antifungal activities, which is unique among structurally similar polyene macrolactams, such as incednine and silvalactam. In this work, we employed different engineering strategies to target glycosylation and acylation biosynthetic machineries within its recently elucidated biosynthetic pathway. Six auroramycin analogs with variations in C-, N-methylation, hydroxylation and extender units incorporation were produced and characterized. By comparing the bioactivity profiles of these analogs, we determined that unique disaccharide motif of auroramycin is essential for its antimicrobial bioactivity. We further demonstrated that C-methylation of the 3, 5-*epi*-lemonose unit, which is unique among structurally similar polyene macrolactams, is key to its antifungal activity.

## Introduction

Natural products (NPs) are an important source for pharmacological applications with a significant proportion of current drugs being natural product or natural product-derived [1]. Advances in genome editing and synthetic biology tools, together with natural product biosynthesis knowledge accumulated over the decades, allows us to better predict, design and build pathways towards the synthesis of natural products [2]. Earlier, we established a rapid and efficient CRISPR-Cas9 strategy for biosynthetic gene cluster (BGC) editing and activation in streptomycetes [3, 4], which opens up opportunities for combinatorial biosynthesis in native streptomycete hosts [5]. Compared to chemical syntheses, combinatorial engineering of native biosynthetic pathways allows us to design and biosynthesize structurally complex chemical analogs without traversing difficult multi-step and possibly low-yielding chemical reactions, thus facilitating elucidation of structure-activity relationships towards an optimized drug lead.

In this study, we describe our efforts to engineer the BGC of antimicrobial compound auroramycin [6, 7] and characterize its structure activity relationship (SAR). Auroramycin (**1**) is a polyene macrolactam that is doubly glycosylated. Sugars are attached in the order of xylosamine and 3, 5-*epi*-lemonose to the polyketide core. Compared to structurally similar natural products (Figure 1), such as the doubly glycosylated incednine [8] and monoglycosylated silvalactam [9], auroramycin is the only polyene macrolactam with reported antifungal activity to date. One of the main structural differences between auroramycin, incednine and silvalactam is their glycosylation pattern (Figure 1). Glycosylation can significantly increase the diversity and complexity of natural products and has often been shown to directly and significantly impact the their bioactivity and pharmacological properties [10, 11]. As such, glycodiversification is an attractive strategy to diversify and optimize bioactivity of NPs. Auroramycin also has an unique isobutyrylmalonyl (ibm) moiety incorporated into its polyene core. This ibm extender unit is relatively rare among NPs [12]. As a mixture of malonyl and methylmalonyl moieties are also incorporated into the polyketide, acyltransferase engineering would also allow us to quickly access new structurally diverse analogs.

**Figure 1.**
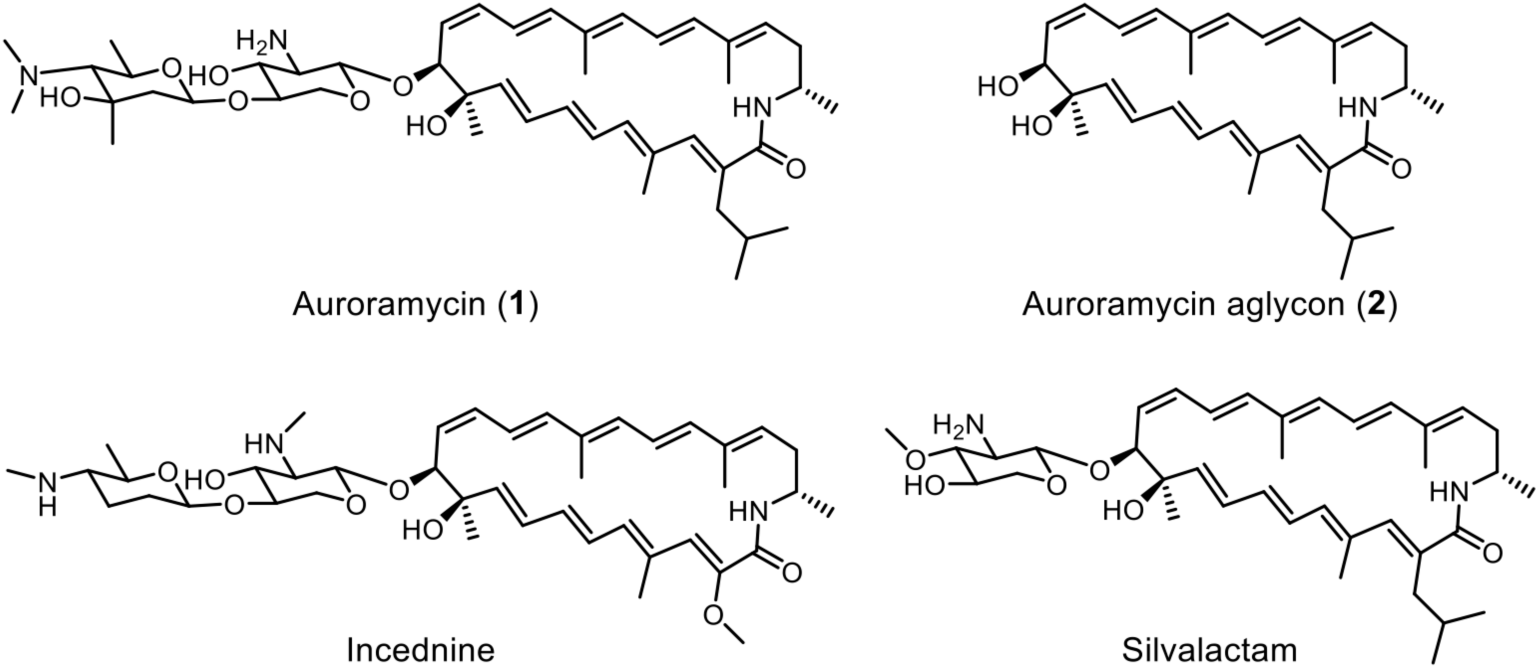
Auroramycin and structurally related polyene macrolactams.

Here, we describe pathway engineering for glycosylation and extender unit incorporation in native *S. roseoporous*. To examine importance of polyketide backbone modifications and its unique disaccharide moiety towards anti-MRSA and antifungal activities, six new auroramycin analogs were produced and characterized.

## Results

### Engineering sugar biosynthesis

We previously proposed the sugar biosynthesis pathways for auroramycin (Figure 2A, [6]). Based on this, deletion of the N,N-dimethyltransferase AurS9 and C-methyltransferase AurS11 should remove N-methylation and C-methylation of 3, 5-*epi*-lemonose to yield compounds **3** and **4** respectively. As expected, **3** and **4** were main products of the respective engineered strains (Figure 2). Production yields of **3** and **4** were 40-80 mg/L, which are comparable to that of auroramycin. The latter observation suggested that the glycosyltransferase for 3, 5-*epi*-lemonose was able to transfer different non-methylated sugars onto the auroramycin scaffold just as efficiently as its cognate sugar substrate.

**Figure 2.**
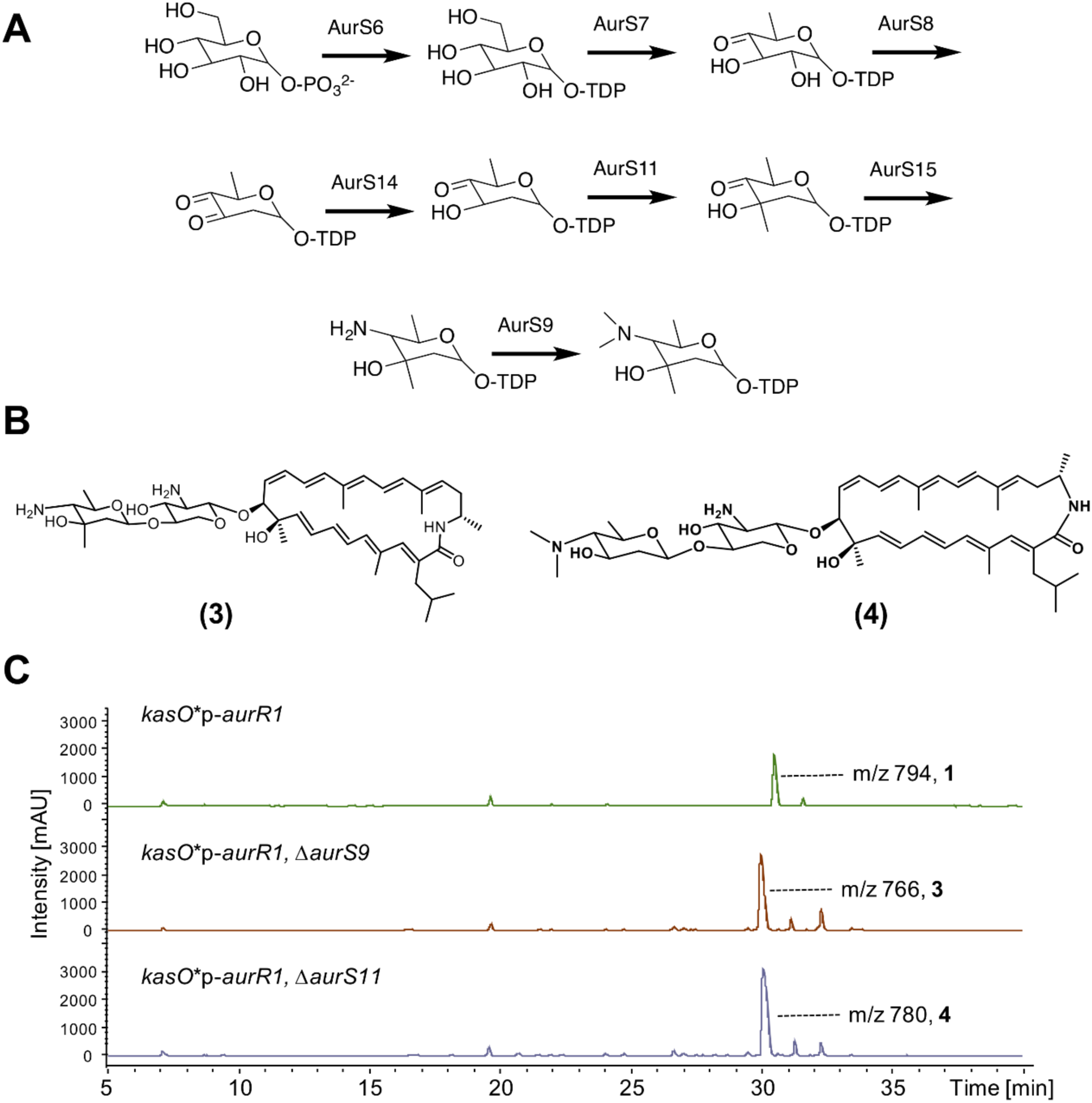
Engineering 3, 5-*epi*-lemonose biosynthesis. (A) Putative sugar biosynthesis pathway. (B) Structures of auroramycin analogs **3** and **4.** (C) Liquid chromatography mass spectrometry (LCMS) spectra of extracts from engineered *S. roseosporus* strains with no modification to native BGC, (top) deletion of *aurS9* and (middle) deletion of *aurS11* (bottom) in the auroramycin BGC.

### Engineering glycosylation

Two glycosylation events take place during auroramycin biosynthesis. However within the BGC, there are four annotated glycosyltransferases (AurS4, S5, S10, S13) in addition to a P450 (AurS12) as its potential auxiliary partner, DesVIII [13, 14]. The four glycosyltransferases have high similarity to DesVII/EryCIII glycosyltransferases which require activation and stabilization by an DesVIII homolog auxiliary partner. A DesVII/DesVIII complex is required for activity. Of these four glycosyltransferases, two were truncated and hypothesized to be inactive; N-terminal helices, hypothesized for DesVIII interactions, and potential sites for substrates and sugar interactions were missing or modified (Figure S9, [13, 14]). Earlier, we obtained aglycon **2** by deleting a 13 kb region in the BGC, which includes the *aurS5, S10, S12, S13* genes [6]. To assign the roles of individual glycosyltransferases and putative auxiliary partner, as well as produce the mono-glycosylated analog **5** for SAR studies, we systematically deleted each of the genes and characterized the products of the engineered *S. roseosporus* strains (Figure 3).

**Figure 3.**
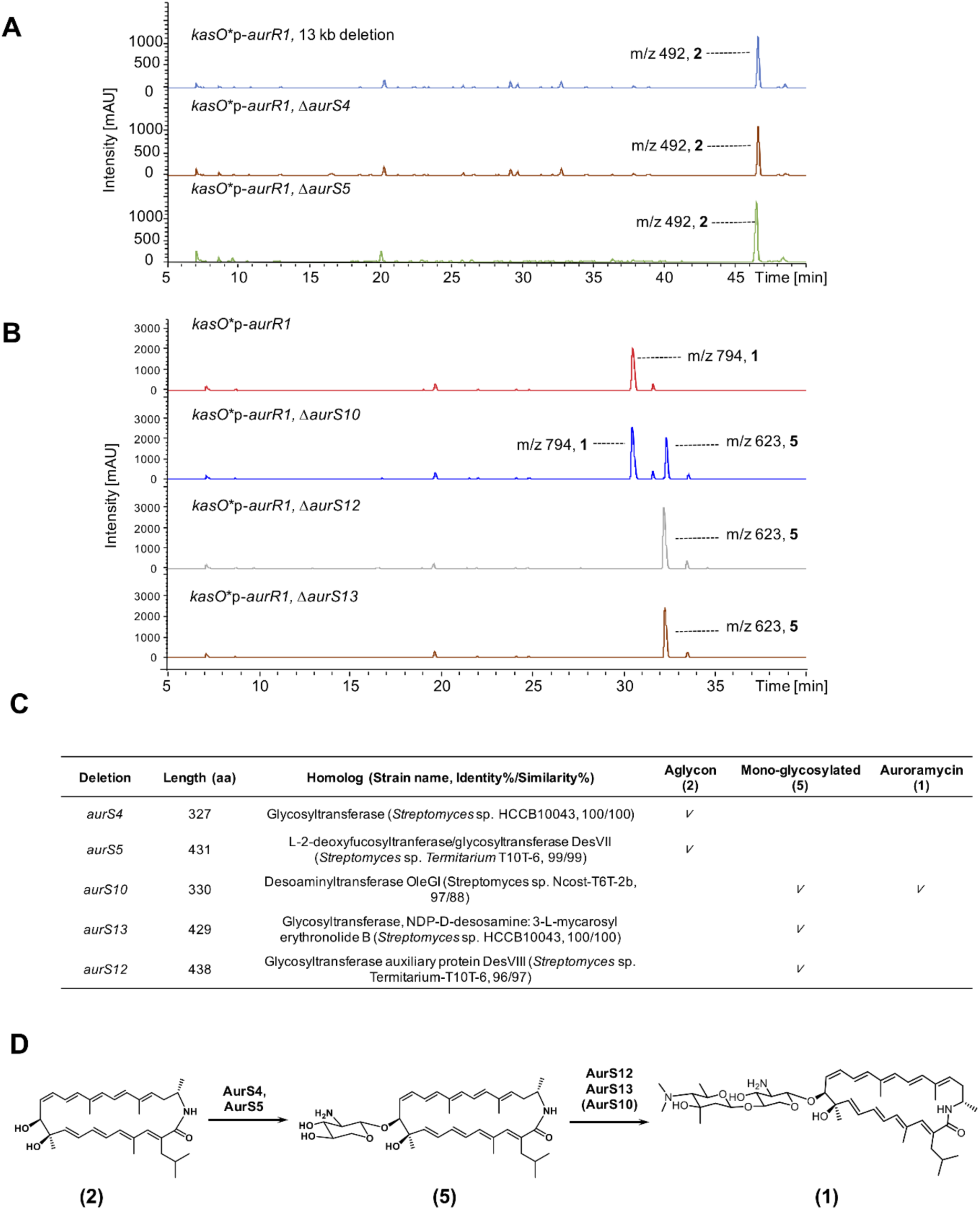
Engineering glycosylation. (A) LCMS spectra of engineered *S. roseosporus* strains with 13 kb and *aurS4* deletions within the BGC. (B) LCMS spectra of engineered *S. roseosporus* strains with no deletion, *aurS10, aurS12* and *aurS13* deletions within the BGC. (C) Table depicting the genes that were deleted, their annotated gene products and resulting metabolite produced by *S. roseosporus* strains carrying the deletion. Length refers to the number of amino acid residues of the indicated gene product. (D) Putative glycosylation scheme of auroramycin. AurS10 is not essential for auroramycin glycosylation but increases the efficiency of the second glycosylation reaction.

Individual deletions of the 5 genes involved in auroramycin glycosylation revealed that different sets and interactions of glycosyltransferases are required for each of the two glycosylation events (Figure 3A-C). The first glycosylation event required the gene products of *aurS4* and *aurS5* as deletion of either gene yielded aglycon **2** (Figure 3A). The second glycosylation event likely involved a more traditional DesVII/VIII complex encoded by *aurS12* and *aurS13*. Purification and characterization of co-expressed AurS12 and AurS13 revealed that AurS13 and AurS12 have similar oligomerization profile as the DesVII/DesVIII complex [13, 14], suggesting that the two proteins form a functional glycosylation complex similar to the latter (Figure S10). Deletion of either *aurS12* and *aurS13* yielded the mono-glycosylated analog **5** (Figure 3B). As aglycon **2** was not observed with *aurS12* deletion, this suggested that the first glycosylation step did not require an auxiliary protein partner for activity. AurS10 was also not essential for auroramycin glycosylation but most likely enhanced the second glycosylation event since deletion of *aurS10* resulted in the production of a mixture of analog **5** and auroramycin (Figure 3B).

Based on these observations, we proposed the following scheme for glycosylation in the auroramycin biosynthetic pathway: xylosamine is first glycosylated by AurS4 and AurS5, after which, 3, 5-*epi*-lemonose is added by AurS12 and AurS13 with AurS10 needed for increased efficiency of the second glycosylation step (Figure 3C, D). Notably, contrary to *in silico* predictions that the truncated gene products of *aurS4* and *aurS10* are non-functional (Figure S9, [15]), deletion of AurS4 and AurS10 has a profound effect on auroramycin’s glycosylation, suggesting a functional role for these truncated gene products.

### Engineering extender unit

To engineer the extender units that constitute the macrocycle core (Figure 1, 4), we first examined a strategy consisting of complementation of an inactivated *cis*-acyltransferase with a *trans*-acting acyltransferase of a different specificity (Figure 4A, [16]). Previous AT complementation examples include complementation of a single module (DEBS Mod6) for the production of 2-desmethyl-6-dEB by malonyltransferase [17] and by a *trans*-acting AT from bryostatin PKS [18]. These studies demonstrated targeted incorporation of malonyl CoA into the polyketide carbon backbone. To engineer the macrocycle core of auroramycin, we chose a highly active malonyl CoA-specific *trans*-acting acyltransferase from the disorazole PKS (Dszs AT, [19]) to functionally rescue inactivated mmCoA/ibmCoA-specific acyltransferases in the auroramycin PKS (Figure 4B).

**Figure 4.**
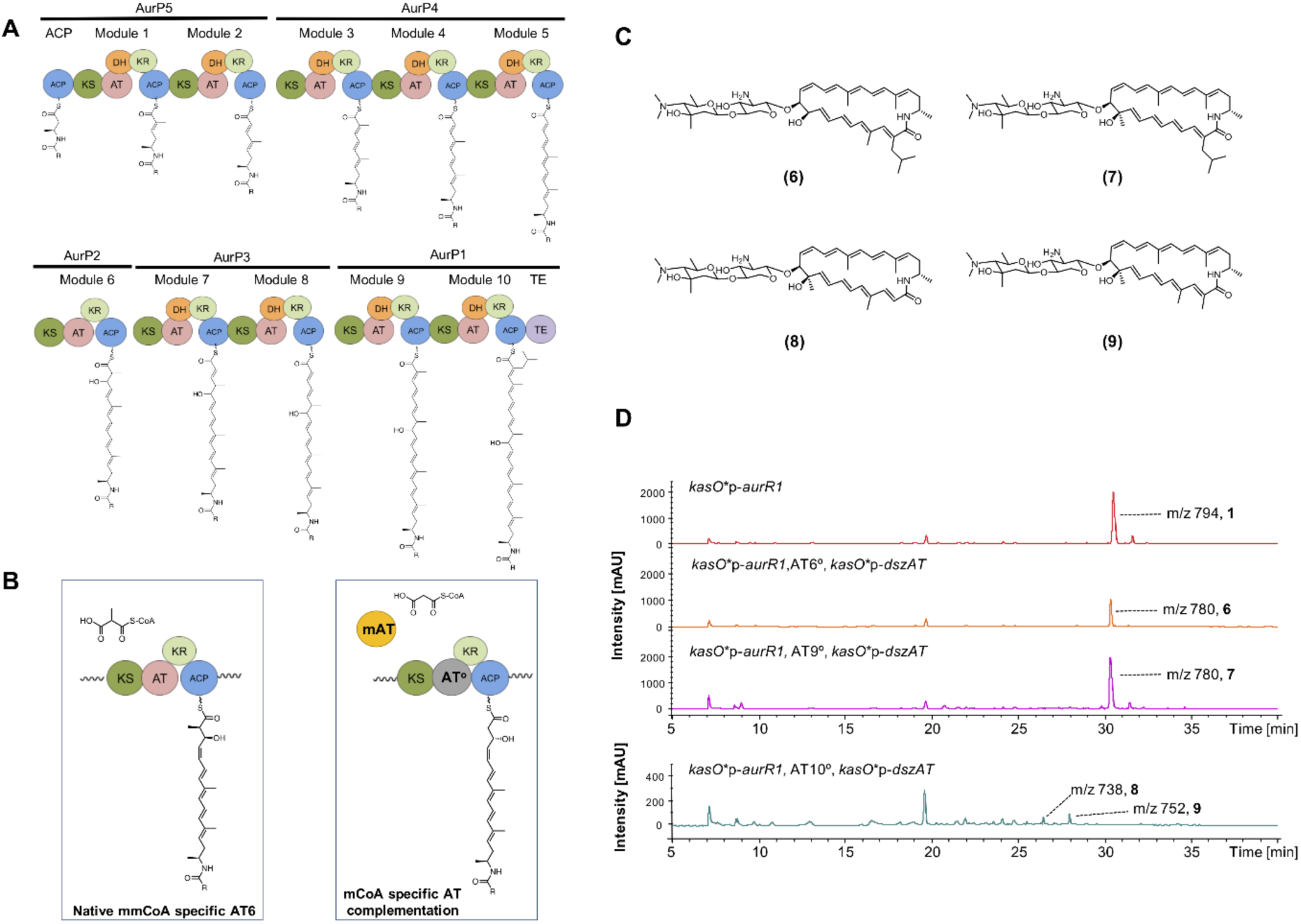
Extender unit engineering of the auroramycin PKS. (A) Polyketide synthase assembly (B) AT complementation schematic (left: native AT in modular PKS, right, complementation of inactivated AT by trans-acting AT) (C) Analogs observed in extender unit engineering (D) LC-MS spectra of the auroramycin-producing strain compared to AT6°, AT9° and AT10° strains complemented by Dszs AT. KS: Ketosynthase, AT: Acyltransferase, KR: Ketoreductase, ACP: Acyl carrier protein, DH: Dehydratase, ER: Enoyl reductase, TE: Thioesterase, mAT: Malonyl CoA-specific acyltransferase.

Dszs AT complementation of inactive AT6 and AT9 (AT6°, AT9°) yielded the expected products **6** and **7** respectively (Figure 4C, D). However, yields differed significantly with **6** produced at 50% yield compared to **7** (Figure 4D). The latter observation was most likely due to substrate specificities of the downstream modules that limited their abilities to process the non-native intermediates [20-22]; the intermediate of AT6° required processing along four modules compared to the AT9° intermediate, which had to be accepted by a single module before cyclization.

With Dszs AT complementation of inactive AT10 (AT10°), instead of obtaining only the malonyl moiety incorporated compound (**8**, Figure 4D), we also found the mmCoA analog **9**. Analogs **8** and **9** were produced at an approximately 1:2 ratio and with yields less than 5% of auroramycin. Substrate preference of Dszs AT for mCoA to mmCoA was observed previously to be approximately more than 46,000-fold [23]. Thus the significant decrease in product yields, along with the observed mmCoA-incorporated product, suggested that C-2 methylation is highly favoured by downstream gatekeeper domains, in particular the thioesterase domain [24]. Due to the low yields of analogs **8** and **9**, their bioactivity was not characterized.

### Post PKS hydroxylation

Among structurally similar natural products such as incednine, silvalactam and auroramycin (Figure 1), post-PKS hydroxylation takes place at a methylated site on the polyene core carbon skeleton. Conservation of this functional group suggests that either hydroxylation or/and methylation at this site might be important for bioactivity. To investigate the role of the additional hydroxyl group at C-10, we examined the production of dehydroxylated analog **10**. In the *aurO1* deletion mutant, only production of the dehydroxylated analog **10** was observed (Figure 5).

**Figure 5.**
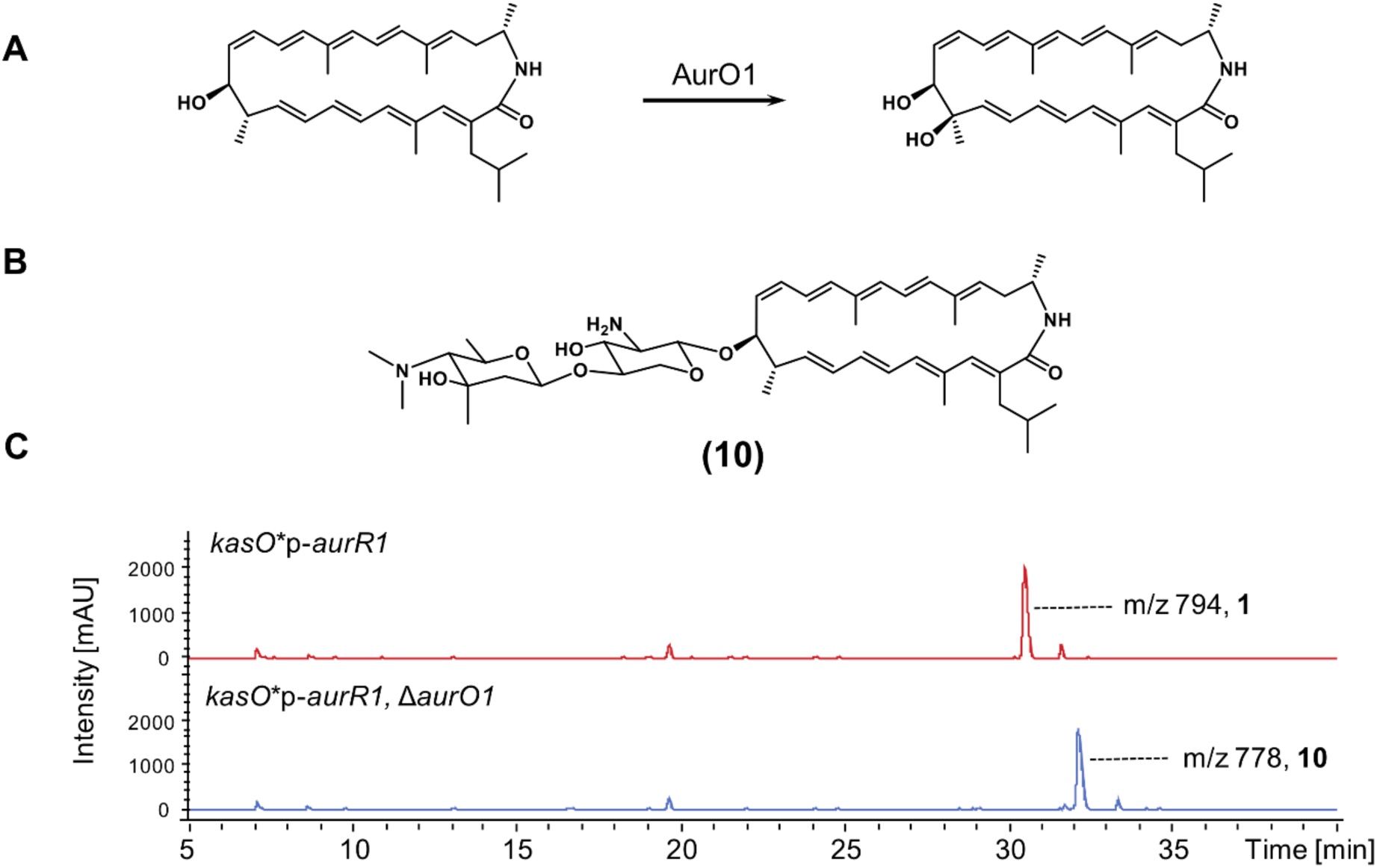
Post PKS hydroxylation. (A) Proposed hydroxylation reaction, (B) the dehydroxylated analog **10**, and (C) respective LC-MS of the engineered *S. roseosporus* in comparison to the auroramycin producing strain.

### Antimicrobial activities of auroramycin analogs

The antifungal activities of the six analogs generated in this study were compared to auroramycin against six fungal and yeast species. Analogs **3** and **4**, which had perturbations in the sugar biosynthesis, gave drastically different results in their bioactivities. The de-*N*-methylated analog **3** retains most of its bioactivity compared to auroramycin. Notably, new bioactivity (against *Kodameae ohmeri*) was observed with analog **3**, suggesting that removing the *N*-methylation on the 3, 5-*epi*-lemonose in auroramycin may improve its potential as a broad spectrum antifungal. In contrast, analog **4** is inactive against the tested fungal and yeast strains, highlighting the importance of the C-methylation on the 3, 5-*epi*-lemonose in auroramycin. As predicted, the mono-glycosylated analog **5**, which is most structurally similar to silvalactam lost all antifungal activity. Our results corroborated previous findings that silvalactam is inactive against yeast *Saccharomyces cerevisiae* [9], and further highlighted the importance of having the additional outer sugar for antifungal bioactivity. Interestingly, analogs **6** and **10**, which contained modifications at C-10 of the macrolactam core, had differing outcomes in their bioactivities. Removal of the hydroxyl group at C-10 (analog **10**) largely retained its bioactivity whereas removal of the methyl group at C-10 (analog **6**) led to complete loss of bioactivity against the tested species. Perturbation of the degree of methylation at other site away from the 3, 5-*epi*-lemonose moiety had little effect on its bioactivity as shown by analog **7**.

**Table 1.** Antifungal activity of auroramycin and its analogs.

|  | Fungi |  |  |  | Yeast |  |
| --- | --- | --- | --- | --- | --- | --- |
| Species | *Candida glabrata* | *Cryptococcus neoformans* | *Candida tropicalis* | *Candida albicans* – azoles resistant | *Saccharomyces cerevisiae* | *Kodameae ohmeri* |
| Strain ID | ATCC 36583 | ATCC 24067 | ATCC 200956 | ATCC 200918 | ATCC 9763 | CDC AR-BANK#0396 |
| Compound | MIC, µg/mL |  |  |  |  |  |
| **1** | 2 | 2 | 1 | 2 | 4 | >128 |
| **3** | 2 | 1 | 1-2 | 2 | 2-4 | 2-4 |
| **4** | >128 | >128 | 64 | >128 | >128 | >128 |
| **5** | >128 | 64 | >128 | >128 | >128 | >128 |
| **6** | >128 | 32 | >128 | >128 | >128 | >128 |
| **7** | 2 | 2 | 2 | 4 | 4 | >128 |
| **10** | 2 | 1 | 2 | 2 | 2 | >128 |

Minimum Inhibitory Concentration (MIC) refers to lowest concentration of an agent that completely inhibits visible growth *in vitro* of the microorganism after 2 days at 28 °C for *Saccharomyces cerevisiae* and *Kodameae ohmeri*, and at 36 °C for *Candida glabrata*, *Cryptococcus neoformans*, *Candida tropicalis* and *Candida albicans*.

We also tested antibacterial activities of the six auroramycin analogs against three Gram-positive bacteria strains, namely methicillin-resistant *Staphylococcus aureus* (MRSA), vancomycin-intermediate methicillin-resistant *Staphylococcus aureus* (VI-MRSA) and vancomycin-resistant *Enterococcus faecalis* (VRE). While there were some notable trends for the antibacterial activities of the analogs, there were distinct differences between the antifungal and antibacterial activities exhibited by the auroramycin analogs. While the analogs generally exhibited either clear retention or complete loss of antifungal bioactivity, alterations to the antibacterial activities for the analogs were more modest. Analog **3** retained its antibacterial activity whereas analog **4** showed some loss of activity, especially against VI-MRSA. This reflected a similar but less drastic effect of the sugar biosynthesis perturbations on antibacterial activity compared to antifungal activity. Even without its outer 3, 5-*epi*-lemonose sugar, analog **5** retained partial antibacterial activity.

While modifications at C-10 of the macrolactam core gave differing outcomes in analogs **6** and **10** antifungal bioactivities, antibacterial bioactivities for analogs **6** and **10** are generally significantly reduced. Similar to analog **4**, analog **7** retained its antifungal activity but incurred a modest 2-4 fold loss in antibacterial activity potency. Like auroramycin, the analogs are inactive against Gram-negative bacteria (MIC >128 μg/mL).

**Table 2.** Antibacterial activity of auroramycin and its analogs.

| Species | *Staphylococcus aureus*<br>(MRSA) | *Staphylococcus aureus*<br>(VI-MRSA) | *Enterococcus faecalis*<br>(VRE) |
| --- | --- | --- | --- |
| Strain ID | Clinical N216 | Clinical Z172 | ATCC51299 |
| Compound | MIC, µg/mL |  |  |
| **1** | 4 | 8 | 2-4 |
| **3** | 4 | 4-8 | 4 |
| **4** | 8-16 | 64 | 8 |
| **5** | 4-8 | 16 | 8 |
| **6** | >128 | >128 | >128 |
| **7** | 8-16 | 16 | 8 |
| **10** | 32 | >128 | 16 |

Minimum Inhibitory Concentration (MIC) refers to lowest concentration of an agent that completely inhibits visible *in vitro* growth of the indicated microorganism after 20 h incubation at 35 °C.

## Discussion

Among structurally similar polyene macrolactams, auroramycin is the only one with reported antifungal activity to date. To identify the functional groups responsible for endowing auroramycin with antifungal and anti-MRSA activities, we sought to engineer auroramycin analogs and characterize the effect of specific chemical structural changes on their bioactivity profiles. Here we targeted sites unique to auroramycin (C, N-methylation on sugars) as well as sites that are common among auroramycin, silvalactam and incednine (unique extender units and hydroxylation). Through rational engineering and CRISPR-Cas mediated genome editing, we could rapidly design, build and test different *S. roseopsorus* mutants. In our study, production yields close to that of auroramycin were achieved for most of the analogs. This could be contributed to minimal disruption to the three-dimensional structure of the polyketide assembly line and sufficiently promiscuous *cis*-acting glycosylation and sugar biosynthesis enzymes. However, there are still engineering bottlenecks, as in the case of swapping the extender unit on C-2 on auroramycin, where minimal product was observed. To access analogs restricted by downstream gatekeeper domains, strategies such as directed evolution or rational design of downstream enzymes to increase substrate promiscuity will have to be examined [25, 26]. However, these strategies are not trivial and should be undertaken for natural products of high interest. Further diversity for targeted functional groups can be achieved by altering enzyme specificities and swapping enzyme domains [16, 27].

By comparing the bioactivity profiles of the six analogs that were achieved in good yields, we were able to evaluate the functional importance of different chemical moieties on auroramycin (Figure 6). From our studies, the unique disaccharide moiety was essential for antifungal and anti-MRSA bioactivity, however anti-MRSA activity can be restored with mono-glycosylation. Interestingly, our results demonstrated unexpected importance of the C-methylation on the outer sugar on bioactivity, especially for antifungal activity. The C-methylation on the outer sugar is unique to auroramycin and could explain auroramycin being the sole antifungal agent among similar polyene macrolactams. Perturbation of the macrolactam core of auroramycin led to mixed outcomes for antifungal activity but mostly resulted in loss of antibacterial activity. Overall, the structural features examined in our study defined auroramycin’s antifungal bioactivity more distinctly than its antibacterial activity. Additional modifications to the sugars and the degree of saturation of the polyene macrolactam core may be explored to improve the antifungal activity of auroramycin.

**Figure 6.**
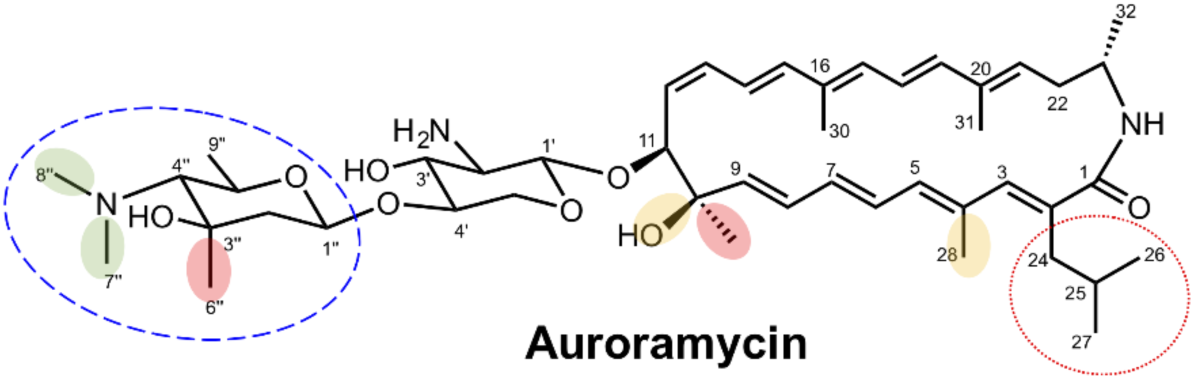
Structural activity map of auroramycin. Red shaded: removal resulted in drastic loss of both antifungal and antibacterial bioactivities; Green shaded: removal resulted in retention of activity or slightly better antifungal activity, including new bioactivity against *K. ohmeri*; Yellow shaded: minimal changes in antifungal activity but loss in potency in antibacterial activity; Dashed blue circle: essential for antifungal activity; enhanced antibacterial potency; Dotted red circle: Due to the low production yields, biological activity of analogs with modification at the ibm moiety were not functionally characterized.

Due to highly efficient, precise and consistent CRISPR-Cas mediated genome editing and advanced DNA assembly methods [28, 29], we were able to accelerate the generation of 12 strains (with at least two genomic edits each) for analog production and screening in this study. Multiplex editing is expected to further accelerate strain construction [3]. Multiplex inactivation of key biosynthetic genes within a target BGC in streptomycetes can also be achieved with base editing using Cas9-deaminase fusions [30]. Last but not least, a major consideration of any genome editing strategy is that it requires the introduction of recombinant DNA, which may be challenging depending on the target actinomycete strain.

## Conclusion

In this work, we used CRISPR-Cas mediated genome editing to accelerate the rational engineering of the antimicrobial auroramycin biosynthetic gene cluster. From a single design-build-test cycle, we were able to make specific changes to the glycosylation pattern and polyene macrolactam core of auroramycin to generate analogs. Six of these analogs were biosynthesized in good yields and their bioactivities were further characterized. By comparing the bioactivities of these analogs, we determined that the unique disaccharide moiety, in particular C-methylation of the outer sugar unit, was important for the antifungal activity of auroramycin.

## Materials and methods

### Growth and conjugation conditions

Unless otherwise indicated, all reagents are obtained from Sigma, St. Louis, USA. Conjugation experiments involving WM3780 *E. coli* strains were performed on R2 agar without sucrose. Unless otherwise indicated, strains are propagated in MGY medium at 30 °C. Spore preparations and conjugation protocols were similar as described before [3]. A typical spore prep contains ∼10^6^–10^7^ spores/mL as determined by serial dilution plating.

### Construction of genome editing plasmids

All DNA manipulations were carried out in OmniMAX™ (Thermo Fisher, Massachusetts, USA). Primers used in this study are listed in Supplementary Table S1. Restriction enzymes were obtained from New England Biolabs. Protospacers were first inserted *via Bbs*I-mediated Golden Gate Assembly before introduction of the respective homology flanks *via* Gibson assembly, as previously described [3]. Detailed description of *aur*S5 deletion strain can also be found in [4].

### Engineering of strains

Precise deletions of individual target genes without affecting intergenic regions were made using a CRISPR-Cas mediated editing strategy ([3], Figure S1-7). To delete *aurS5*, we had to use a different Cas protein [4]. As described elsewhere [6], in order to activate the entire BGC for production of the corresponding analogs, *luxR* in these edited strains was also placed under a strong constitutive *kasO*\* promoter [31].

AT domains in modules 6, 9 and 10 were independently inactivated with a single active site serine to alanine mutation (Figure S8). A gene cassette of Dszs AT, placed under *kasO\**p, was integrated into the *attB* site of the genome [32]. The *luxR* gene within the BGC in these strains was also under *kasO\**p to activate its production.

### Validation of promoter knock-in and genome editing

Genomic DNA from wild type and exconjugants from the indicated strains were isolated from liquid cultures using the Blood and Tissue DNeasy kit (Qiagen, Hilden, Germany) after pretreating the cells with 20 mg/mL lysozyme for 0.5–1 h at 30 °C. PCR was performed using control primers beyond the homology regions with KODXtreme Taq polymerase (Millipore, Massachusetts, USA). Where indicated, PCR products were subjected to digest with specific restriction enzymes to differentiate between PCR products of wild type genomic sequences and successful genome editing by knock-ins. Positive samples were purified using ExoSAP-IT™ (Affymetrix USB, Massachusetts, USA) and validated by Sanger sequencing.

### Fermentation

Fermentation of analogs and auroramycin was performed as described elsewhere [6] using the indicated engineered *Streptomyces roseosporus* NRRL 15998 strains.

### Isolation and characterization of analogs

#### General considerations

Optical rotation was obtained on a JASCO P1030 polarimeter using a micro-cell (light path 1 cm). IR spectrum was taken on a PerkinElmer Spectrum 100 FT-IR spectrometer. UV spectra were recorded on Shimadzu UV-2450 UV-Vis spectrophotometer. CD spectra were taken on a JASCO J-810 CD Spectropolarimeter. HRMS spectra were measured on an Agilent 6545 Quadrupole-Time-of-Flight (Q-TOF) mass spectrometer. ^1^H, ^13^C, and 2D-NMR spectra were recorded on a Bruker 400 MHz and Varian VNMRS 700 MHz spectrometers and calibrated using residual non-deuterated solvent (CD_2_Cl_2_: *δ*H = 5.32 ppm, *δ*C = 53.84 ppm) as an internal reference.

#### Analog 3

To the crude ethyl acetate extracts from 200 agar fermentation plates (∼6 L), cold ethyl acetate (20 mL x 2) was added, sonicated for 1-2 minutes and centrifuged to separate the yellow solution and yellow solid. Cold methanol (10 mL) was then added to the yellow solid, sonicated and centrifuged to separate the yellow solution and analog **3** was obtained as a white/pale yellow solid (238.0 mg). [α]_D_^25^: –124.6° (*c* 0.09, 1:1 CH_2_Cl_2_:MeOH); see Figure S11 for structural assignment; IR (KBr): *ν* = 3353, 2960, 2925, 2857, 1744, 1628, 1577, 1526, 1459, 1385, 1236, 1099, 1077, 985, 962, 881, 847 cm^−1^; UV/VIS (DMSO): *λ*max (log *ε*) = 322.5 (2.62) nm; HRMS *m/z*: 794.5000 [(M+H)^+^ calcd. for C_44_H_68_N_3_O_8_, 766.5006].

#### Analog 4

To the crude ethyl acetate extracts from 125 agar fermentation plates (∼4 L), cold ethyl acetate (20 mL x 2) was added, sonicated for 1-2 minutes and centrifuged to separate the yellow solution and yellow solid. Cold methanol (15 mL) was then added to the yellow solid, sonicated and centrifuged to separate the yellow solution and analog **4** was obtained as a white/pale yellow solid (315.0 mg). [α]_D_^25^: –301.6° (*c* 0.25, 1:1 CH_2_Cl_2_:MeOH); ^1^H and ^13^C NMR: see Table S2; IR (KBr): *ν* = 3353, 2953, 2925, 2871, 1630, 1577, 1525, 1459, 1384, 1284, 1234, 1167, 1129, 1079, 1058, 985, 960 cm^−1^; UV/VIS (1:1 CH_2_Cl_2_:MeOH): *λ*max (log *ε*) = 317.5 (1.25) nm; HRMS *m/z*: 780.5161 [(M+H)^+^ calcd. for C_45_H_70_N_3_O_8_, 780.5163].

#### Analog 5

To the crude ethyl acetate extracts from 184 agar fermentation plates (∼6 L), cold ethyl acetate (30 mL) was added, sonicated for 1-2 minutes and centrifuged to separate the yellow solution and yellow solid. Cold methanol (5 mL) was then added to the yellow solid, sonicated and centrifuged to separate the yellow solution and analog **5** was obtained as a white/pale yellow solid (92.1 mg). [α]_D_^25^: –233.5° (*c* 0.2, 1:1 CH_2_Cl_2_:MeOH); ^1^H and ^13^C NMR: see Table S2; IR (KBr): *ν* = 3350, 2925, 2854, 1712, 1628, 1530, 1463, 1386, 1285, 1147, 1077, 1057, 985, 961 cm^−1^; UV/VIS (MeOH): *λ*max (log *ε*) = 317.5 (1.74) nm; HRMS *m/z*: 623.4056 [(M+H)^+^ calcd. for C_37_H_55_N_2_O_6_, 623.4060].

#### Analog 6

To the crude ethyl acetate extracts from 200 agar fermentation plates (∼4 L), cold diethyl ether (10 mL x 2) was added, sonicated for 1-2 minutes and centrifuged to separate the yellow solution and R22-AT6 was obtained as a brown solid (107 mg). [α]_D_^25^: –235.1° (*c* 0.11, 1:1 CH_2_Cl_2_:MeOH); see Figure S12 for structural assignment; IR (KBr): *ν* = 3333, 2953, 2924, 2852, 1665, 1580, 1465, 1385, 1304, 1270, 1238, 1160, 1096, 1029, 983, 960, 881 cm^−1^; UV/VIS (DMSO): *λ*max (log *ε*) = 320 (2.30) nm; HRMS *m/z*: 780.5156 [(M+H)^+^ calcd. for C_45_H_69_N_3_O_8_ 780.5157].

#### Analog 7

To the crude ethyl acetate extracts from 191 agar fermentation plates (∼4 L), cold diethyl ether (10 mL x 2) was added, sonicated for 1-2 minutes and centrifuged to separate the yellow solution and R22-AT9 was obtained as a brown solid (165 mg). [α]_D_^25^: 51.8° (*c* 0.13, 1:1 CH_2_Cl_2_:MeOH); see Figure S13 for structural assignment; IR (KBr): *ν* = 3356, 2953, 2925, 2871, 1668, 1641, 1609, 1518, 1456, 1377, 1270, 1237, 1161, 1145, 1094, 1058, 995 cm^−1^; UV/VIS (DMSO): *λ*max (log *ε*) = 328.5 (2.57) nm; HRMS *m/z*: 780.5161 [(M+H)^+^ calcd. for C_45_H69N_3_O_8_, 780.5157].

#### Analog 10

To the crude ethyl acetate extracts from 180 agar fermentation plates (∼6 L), cold Et_2_O (10-15 mL x 2) was added, sonicated for 1-2 minutes and centrifuged to separate the yellow solution and brown solid. The brown solid was washed with cold 5:1 Et_2_O:acetone (2 mL x 3) and analog **10** was obtained as a brown solid (69.7 mg). [α]_D_^25^: –261.9° (*c* 0.25, 1:1 CH_2_Cl_2_:MeOH); ^1^H and ^13^C NMR: see Table S3; IR (KBr): *ν* = 3333, 2926, 2867, 1661, 1515, 1456, 1382, 1274, 1238, 1156, 1097, 1058, 988, 962 cm^−1^; UV/VIS (MeOH): *λ*max (log *ε*) = 317.5 (1.90) nm; HRMS *m/z*: 778.5361 [(M+H)^+^ calcd. for C_46_H_72_N_3_O_7_, 778.5370].

### Antifungal assays

Measurements against indicated fungal strains were conducted at Eurofins Panlabs Taiwan (www.eurofins.com/epds), according to methodology described by the Clinical and Laboratory Standards Institute (CLSI) (M38-A, M27-A2).

### Bacterial assays

Minimum inhibition concentration (MIC) values were determined by the broth microdilution method, as recommended by the Clinical and Laboratory Standards Institute with slight modifications. Briefly, purified auroramycin analogs were dissolved in DMSO, then diluted in Mueller-Hinton broth containing 0.2% DMSO. The organisms were tested at 5 x 10^5^ CFU/mL. The MICs were read at 20 h after 35 °C incubation.

## List of abbreviations

SAR: Structure activity relationship
Ibm: Isobutyrylmalonyl
Dszs: Disorazole synthase
PKS: Polyketide synthase
KS: Ketosynthase
AT: Acyltransferase
KR: Ketoreductase
ACP: Acyl carrier protein
DH: Dehydratase
ER: Enoyl reductase
TE: Thioesterase
mAT: malonyl CoA-specific acyltransferase.
MRSA: Methicillin-resistant *Staphylococcus aureus*
HRMS: High resolution mass spectrometry
LCMS: Liquid chromatography mass spectrometry

## Declarations

### Ethics approval and consent to participate

not applicable

### Consent for publication

not applicable

### Availability of data and material

All data generated or analysed during this study are included in this published article [and its supplementary information files].

### Competing interests

The authors declare that they have no competing interests

### Funding

This work is supported by the Agency for Science, Technology and Research (A*STAR), Singapore and National Research Foundation, Singapore [NRF2013-THE001-094 to MMZ, FTW, YHL] and the A*STAR Visiting Investigator Program [HZ].

## Authors’ contributions

WLY performed and analysed conjugation and fermentation experiments.

EH and LLT constructed plasmids for genetic manipulations.

YWL, KCC, YHL performed chemical purifications and characterizations.

D-JT, YWJ, T-LL, K-SS performed and analysed bacterial assays.

HZ contributed to idea conception.

ELA supervised WLY.

MMZ contributed to idea conception and project design.

YHL and FTW conceived project design, analysed data, prepared and revised the manuscript.

All authors read and approved the final manuscript.

## Acknowledgements

The authors gratefully acknowledge the late Sydney Brenner for his discussion and support.

## References

1. Newman DJ, Cragg GM. Natural products as sources of new drugs from 1981 to 2014. Journal of natural products. 2016 Feb 7;79(3):629–61.

2. Zhang MM, Qiao Y, Ang EL, Zhao H. Using natural products for drug discovery: the impact of the genomics era. Expert opinion on drug discovery. 2017 May 4;12(5):475–87.

3. Zhang MM, Wong FT, Wang Y, Luo S, Lim YH, Heng E, Yeo WL, Cobb RE, Enghiad B, Ang EL, Zhao H. CRISPR–Cas9 strategy for activation of silent *Streptomyces* biosynthetic gene clusters. Nature chemical biology. 2017 Jun;13(6):607.

4. Yeo WL, Heng E, Tan LL, Lim YW, Lim YH, Hoon S, Zhao H, Zhang MM, Wong FT. Characterization of Cas proteins for CRISPR-Cas editing in streptomycetes. Biotechnology and Bioengineering. 2019 May 15

5. Weissman KJ, Leadlay PF. Combinatorial biosynthesis of reduced polyketides. Nature reviews microbiology. 2005 Dec;3(12):925.

6. Lim YH, Wong FT, Yeo WL, Ching KC, Lim YW, Heng E, Chen S, Tsai DJ, Lauderdale TL, Shia KS, Ho YS, Hoon S, Ang EL, Zhang MM, Zhao H. Auroramycin: A Potent Antibiotic from *Streptomyces roseosporus* by CRISPR-Cas9 Activation. ChemBioChem. 2018 Aug 16;19(16):1716–9.

7. Wong JH, Alfatah M, Kong KW, Hoon S, Yeo WL, Ching KC, Goh CJ, Zhang MM, Lim YH, Wong FT, Arumugam P. Chemogenomic profiling in yeast reveals antifungal mode-of-action of polyene macrolactam auroramycin. PloS one. 2019 Jun 10;14(6):e0218189.

8. Futamura Y, Sawa R, Umezawa Y, Igarashi M, Nakamura H, Hasegawa K, Yamasaki M, Tashiro E, Takahashi Y, Akamatsu Y, Imoto M. Discovery of incednine as a potent modulator of the anti-apoptotic function of Bcl-xL from microbial origin. Journal of the American Chemical Society. 2008 Feb 13;130(6):1822–3.

9. Schulz D, Nachtigall J, Geisen U, Kalthoff H, Imhoff JF, Fiedler HP, Süssmuth RD. Silvalactam, a 24-membered macrolactam antibiotic produced by *Streptomyces* sp. Tü 6392. The Journal of antibiotics. 2012 Jul;65(7):369.

10. Huang G, Lv M, Hu J, Huang K, Xu H. Glycosylation and activities of natural products. Mini reviews in medicinal chemistry. 2016 Aug 1;16(12):1013–6.

11. Elshahawi SI, Shaaban KA, Kharel MK, Thorson JS. A comprehensive review of glycosylated bacterial natural products. Chemical Society Reviews. 2015;44(21):7591–697.

12. Lechner A, Wilson MC, Ban YH, Hwang JY, Yoon YJ, Moore BS. Designed biosynthesis of 36-methyl-FK506 by polyketide precursor pathway engineering. ACS synthetic biology. 2012 Nov 5;2(7):379–83.

13. Moncrieffe MC, Fernandez MJ, Spiteller D, Matsumura H, Gay NJ, Luisi BF, Leadlay PF. Structure of the glycosyltransferase EryCIII in complex with its activating P450 homologue EryCII. Journal of molecular biology. 2012 Jan 6;415(1):92–101.

14. Borisova SA, Liu HW. Characterization of glycosyltransferase DesVII and its auxiliary partner protein DesVIII in the methymycin/pikromycin biosynthetic pathway. Biochemistry. 2010 Aug 24;49(37):8071–84.

15. Takaishi M, Kudo F, Eguchi T. Identification of the incednine biosynthetic gene cluster: characterization of novel β-glutamate-β-decarboxylase IdnL3. The Journal of antibiotics. 2013 Dec;66(12):691.

16. Musiol-Kroll E, Wohlleben W. Acyltransferases as tools for polyketide synthase engineering. Antibiotics. 2018 Sep;7(3):62.

17. Kumar P, Koppisch AT, Cane DE, Khosla C. Enhancing the modularity of the modular polyketide synthases: transacylation in modular polyketide synthases catalyzed by malonyl-CoA: ACP transacylase. Journal of the American Chemical Society. 2003 Oct 31;125(47):14307–12.

18. Lopanik NB, Shields JA, Buchholz TJ, Rath CM, Hothersall J, Haygood MG, Håkansson K, Thomas CM, Sherman DH. In vivo and in vitro trans-acylation by BryP, the putative bryostatin pathway acyltransferase derived from an uncultured marine symbiont. Chemistry & biology. 2008 Nov 24;15(11):1175–86.

19. Wong FT, Jin X, Mathews II, Cane DE, Khosla C. Structure and mechanism of the trans-acting acyltransferase from the disorazole synthase. Biochemistry. 2011 Jul 5;50(30):6539–48.

20. Jenner M, Frank S, Kampa A, Kohlhaas C, Pöplau P, Briggs GS, Piel J, Oldham NJ. Substrate specificity in ketosynthase domains from trans-AT polyketide synthases. Angewandte Chemie International Edition. 2013 Jan 21;52(4):1143–7.

21. Murphy AC, Hong H, Vance S, Broadhurst RW, Leadlay PF. Broadening substrate specificity of a chain-extending ketosynthase through a single active-site mutation. Chemical Communications. 2016;52(54):8373–6.

22. Zhang L, Hashimoto T, Qin B, Hashimoto J, Kozone I, Kawahara T, Okada M, Awakawa T, Ito T, Asakawa Y, Ueki M. Characterization of giant modular PKSs provides insight into genetic mechanism for structural diversification of aminopolyol polyketides. Angewandte Chemie International Edition. 2017 Feb 6;56(7):1740–5.

23. Dunn BJ, Watts KR, Robbins T, Cane DE, Khosla C. Comparative analysis of the substrate specificity of trans-versus cis-acyltransferases of assembly line polyketide synthases. Biochemistry. 2014 Jun 9;53(23):3796–806.

24. McMahon MD, Prather KL. Functional screening and in vitro analysis reveal thioesterases with enhanced substrate specificity profiles that improve short-chain fatty acid production in Escherichia coli. Appl. Environ. Microbiol. 2014 Feb 1;80(3):1042–50.

25. Zabala AO, Cacho RA, Tang Y. Protein engineering towards natural product synthesis and diversification. Journal of industrial microbiology & biotechnology. 2012 Feb 1;39(2):227–41.

26. Twigg F, Skyrud D, Li J, Zhang W. Engineering Enzymes for Natural Product Biosynthesis and Diversification. InModern Biocatalysis 2018 May 10 (pp. 261–286).

27. Bernhardt P, O’Connor SE. Opportunities for enzyme engineering in natural product biosynthesis. Current opinion in chemical biology. 2009 Feb 1;13(1):35–42.

28. Engler C, Gruetzner R, Kandzia R, Marillonnet S. Golden gate shuffling: a one-pot DNA shuffling method based on type IIs restriction enzymes. PloS one. 2009 May 14;4(5):e5553.

29. Gibson DG, Young L, Chuang RY, Venter JC, Hutchison III CA, Smith HO. Enzymatic assembly of DNA molecules up to several hundred kilobases. Nature methods. 2009 May;6(5):343.

30. Zhong Z, Guo J, Deng L, Chen L, Wang J, Li S, Xu W, Deng Z, Sun Y. Base editing in *Streptomyces* with Cas9-deaminase fusions. bioRxiv. 2019 Jan 1:630137.

31. Wang W, Li X, Wang J, Xiang S, Feng X, Yang K. An engineered strong promoter for streptomycetes. Appl. Environ. Microbiol.. 2013 Jul 15;79(14):4484–92.

32. Combes P, Till R, Bee S, Smith MC. The *Streptomyces* genome contains multiple pseudo-attB sites for the φC31-encoded site-specific recombination system. Journal of bacteriology. 2002 Oct 15;184(20):5746–52.

