## Supplemental data for "Biosynthetic engineering of the antifungal, anti-MRSA auroramycin"

**Supplemental Table S1**. Oligonucleotides used in this study

- Available as “Oligo_Jan.xls”

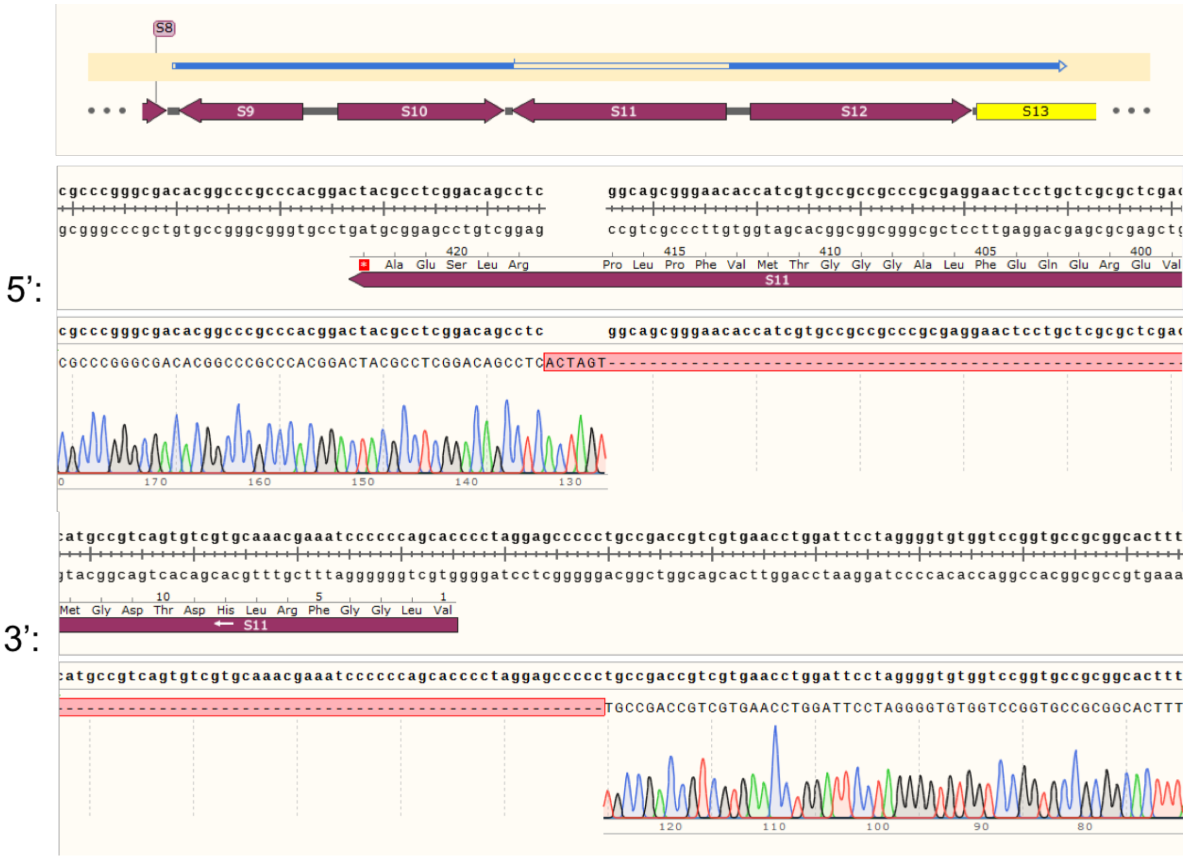


**Figure S1.** Sanger sequencing of *aurS9* deletion mutant.

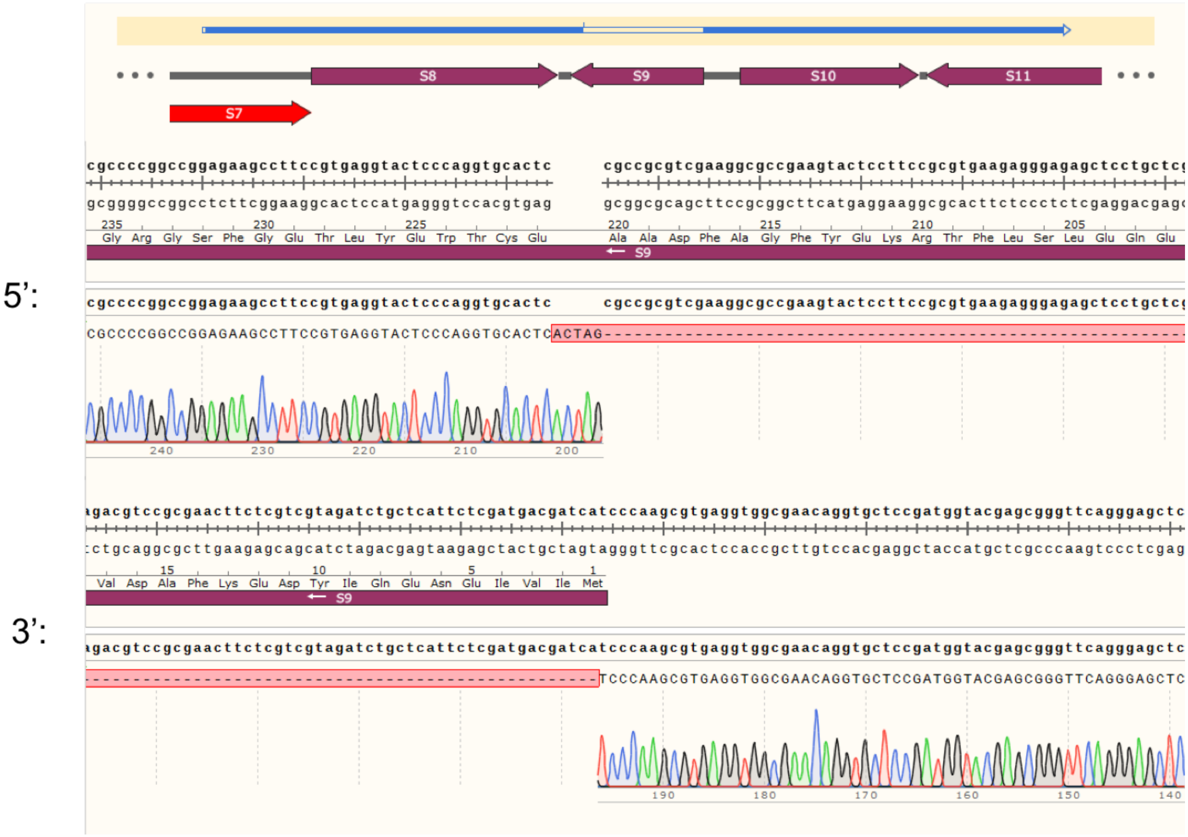


**Figure S2.** Sanger sequencing of *aurS11* deletion mutant.

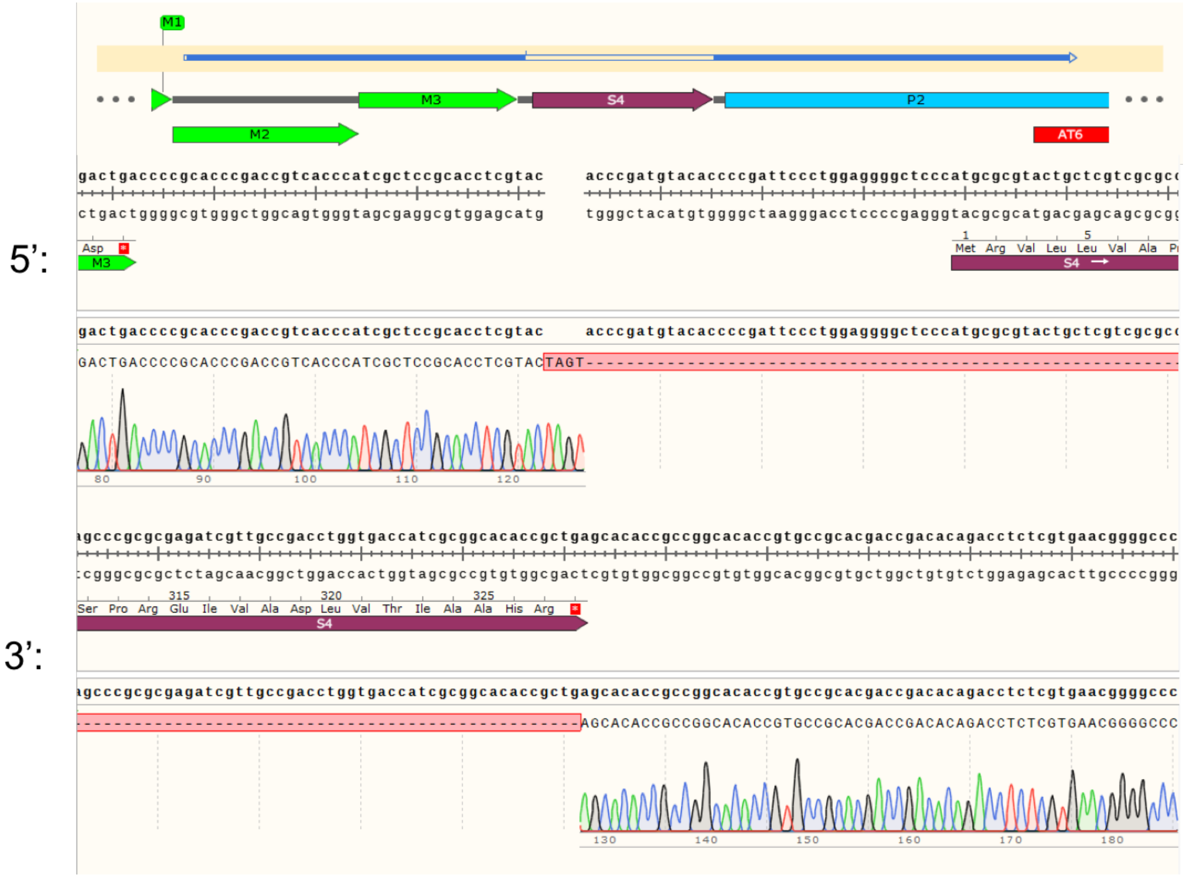


**Figure S3.** Sanger sequencing of *aurS4* deletion mutant.

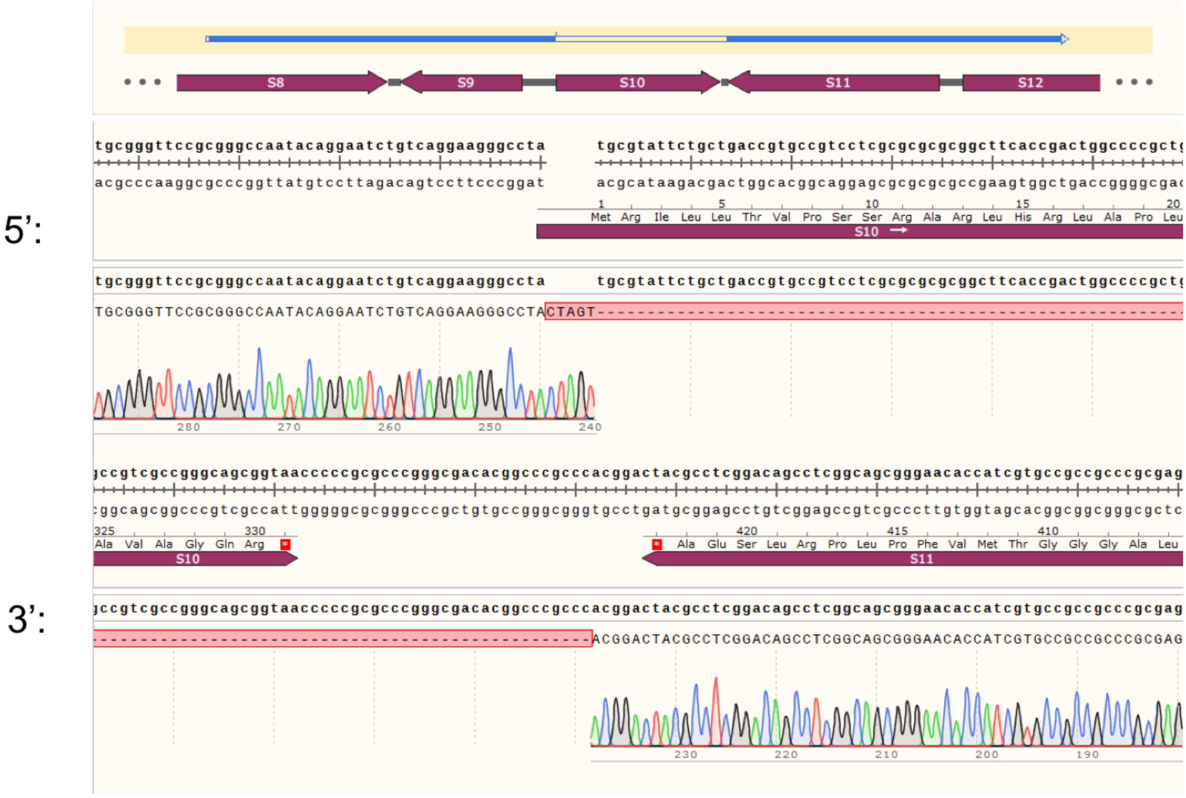


**Figure S4.** Sanger sequencing of *aurS10* deletion mutant.

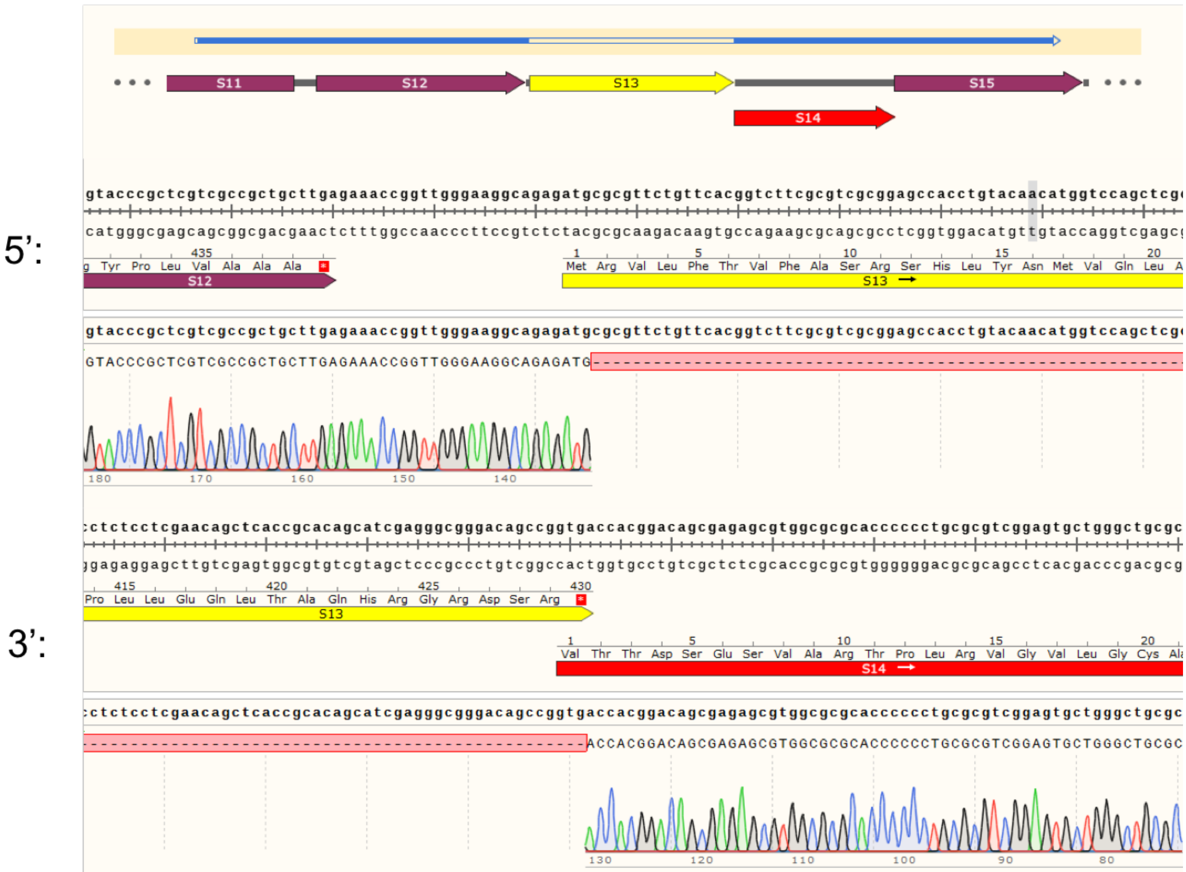


**Figure S5**. Sanger sequencing of *aurS12* deletion mutant.

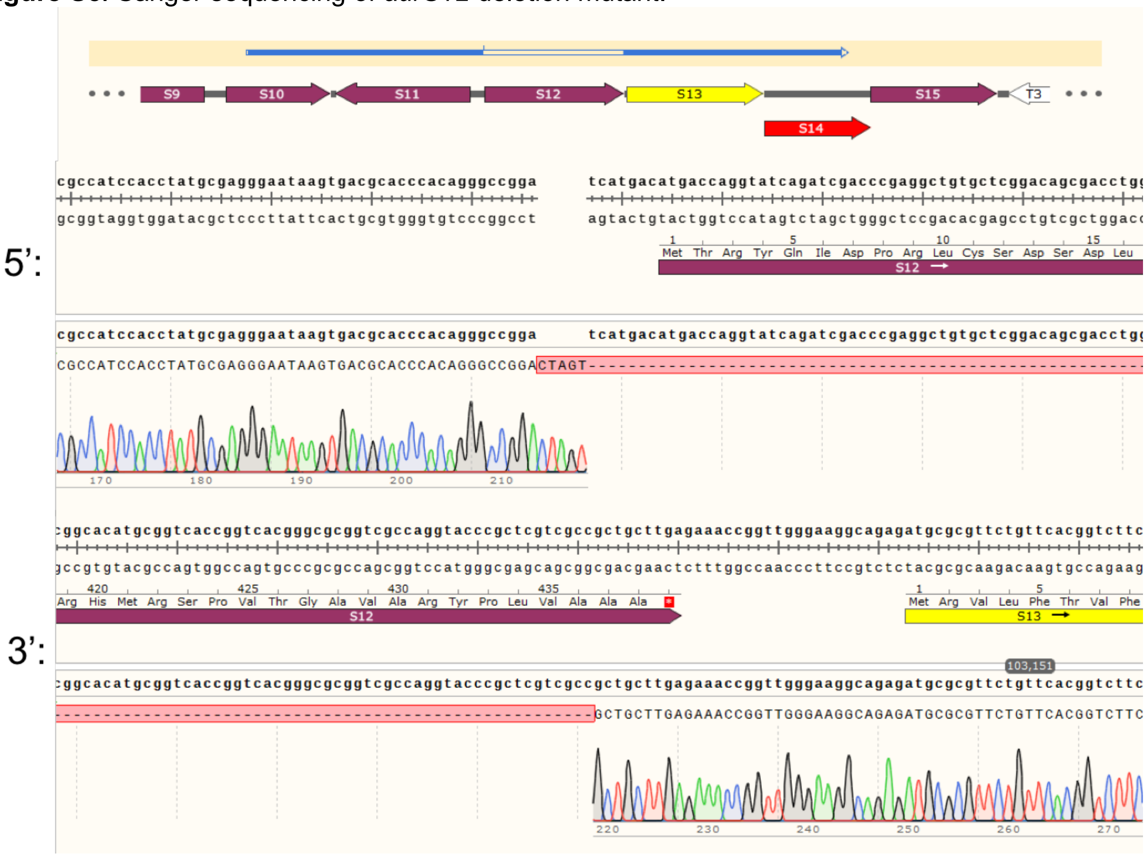


**Figure S6.** Sanger sequencing of *aurS13* deletion mutant.

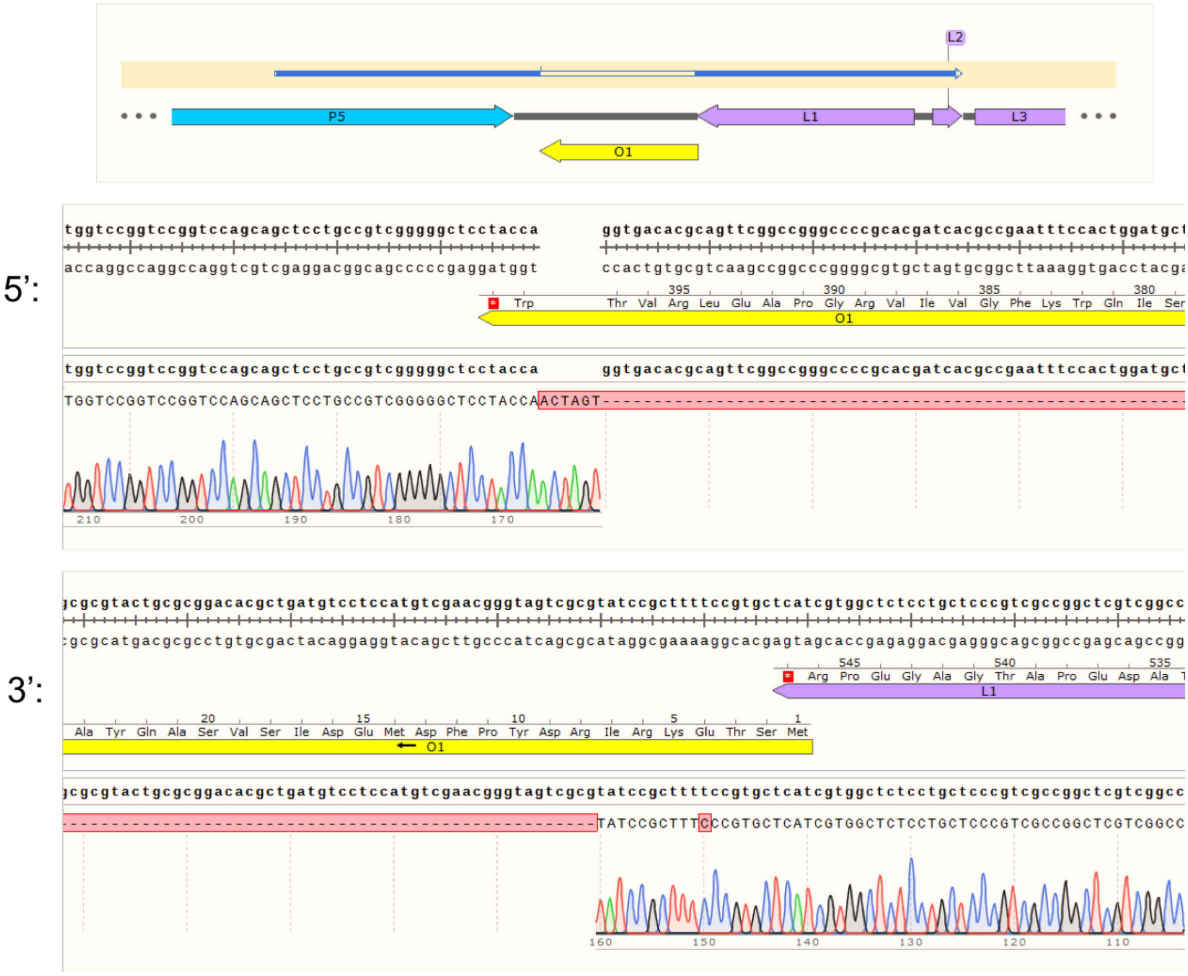


**Figure S7.** Sanger sequencing of *aurO1Δ* mutant. A *Spe*I site was inserted in place of the *aurO1* deletion. A single base mutation was also observed on the remaining segment of *aurO1*, however we predict minimal effects of this single base mutation on our experiments.

.

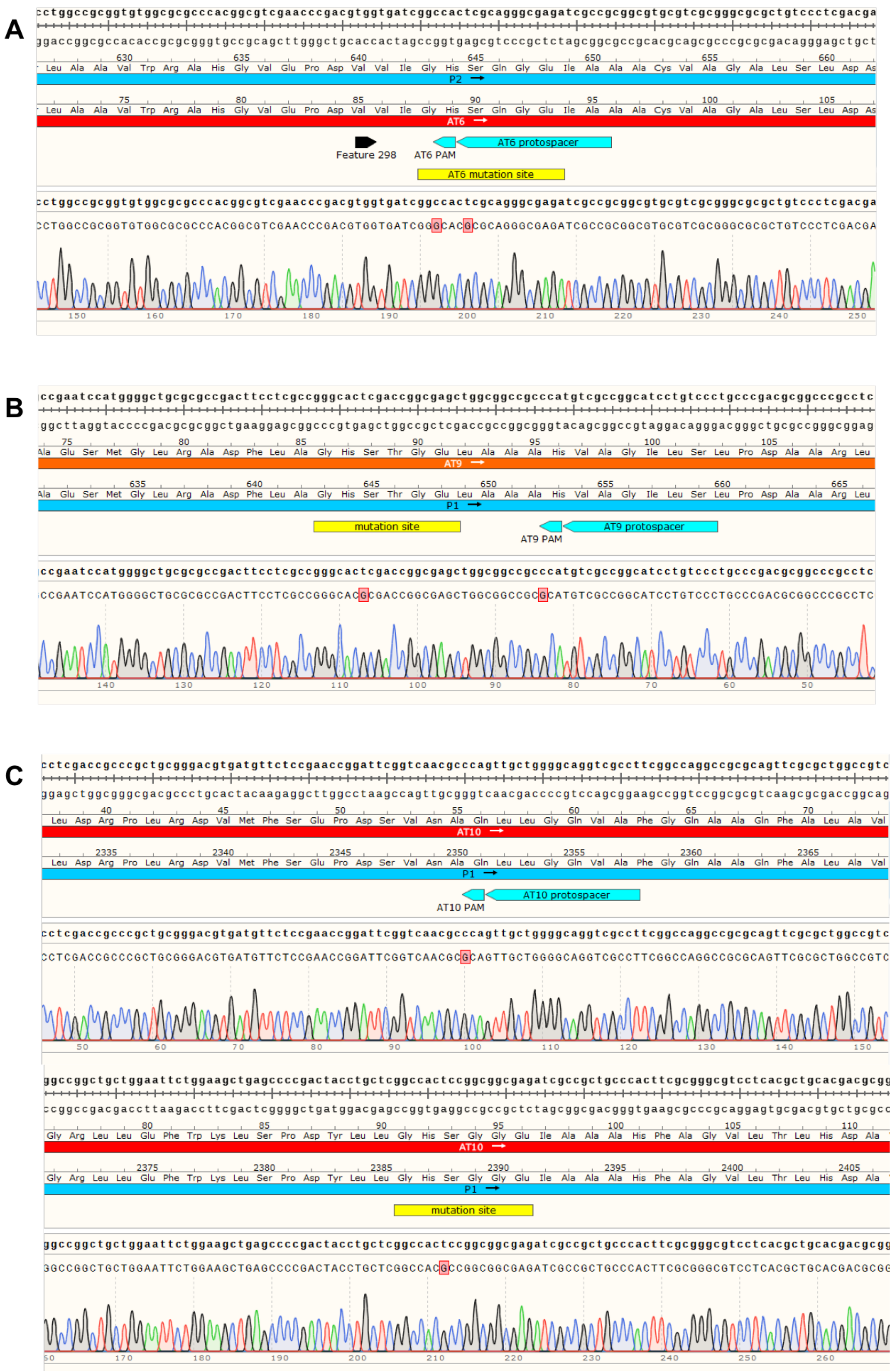


**Figure S8.** Sanger sequencing of AT mutants. Active site conserved serine to alanine mutants for (A) AT6 (B) AT9 and (C) AT10. PAM sites were also mutated to stop re-cutting by CRISPR-Cas.

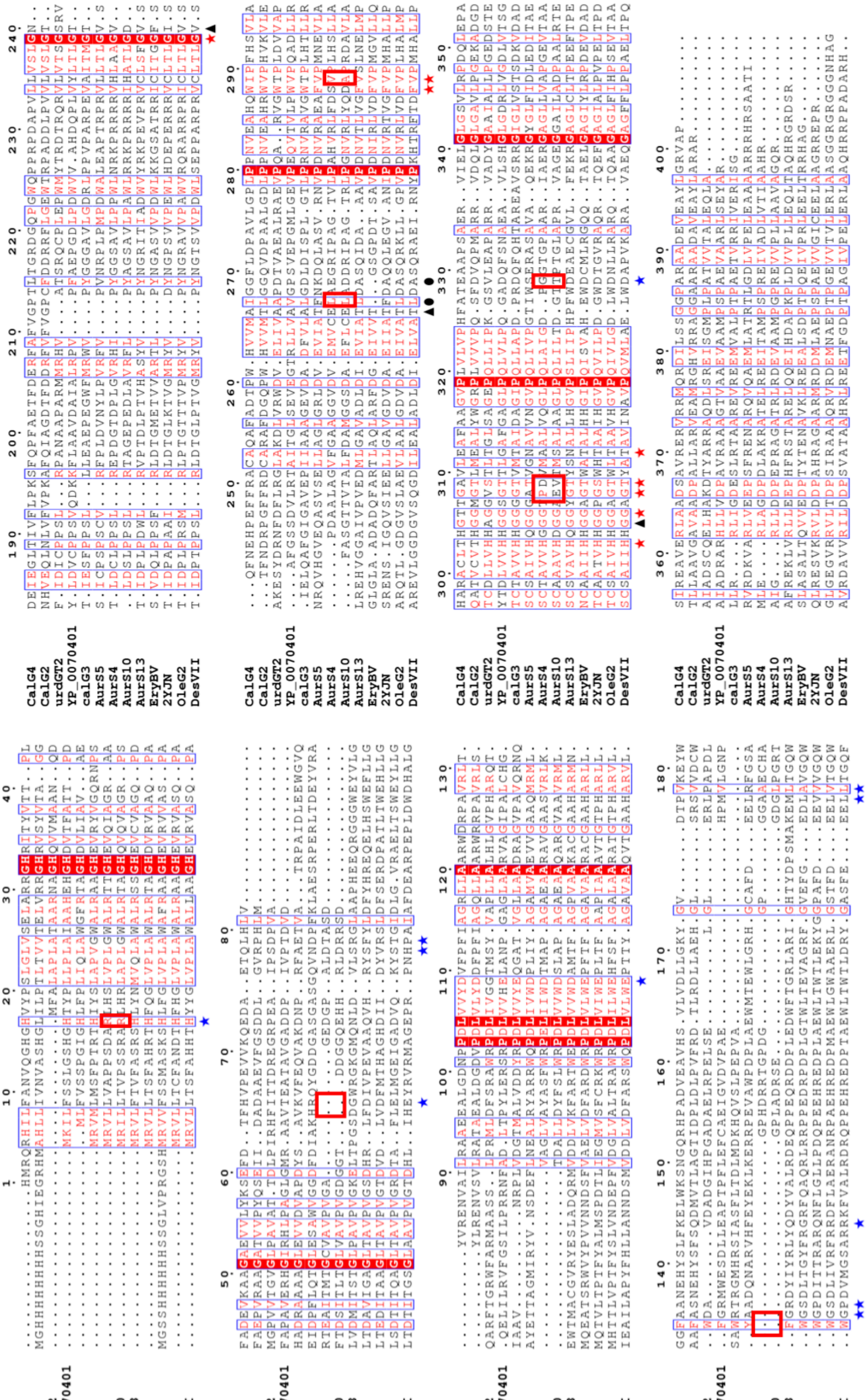

**Figure S9 Alignment of glycosyltranferases.** Putative residues, based on EryCIII (2YJN, Moncrieffe et al, 2012) involved in acceptor and donor nucleotide binding are highlighted. Residues marked with blue stars are involved in acceptor binding while residues marked with red stars are involved in both TDP and UDP binding. Black triangles and circles denote residues that are involved in only UDP or TDP binding. Distinct differences within the truncated glycosyltransferases for proposed residues participating in acceptor or sugar binding are boxed in red.

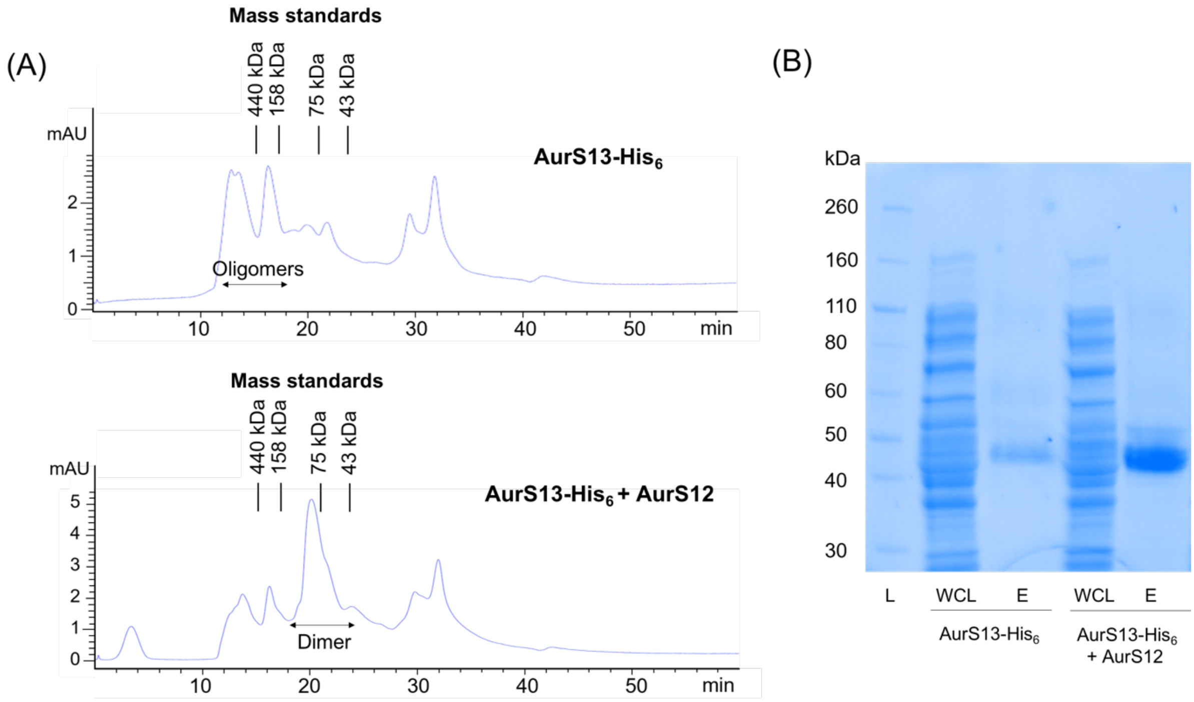


**Figure S10** Purification and characterization of His-tagged AurS13 expressed alone or His-tagged AurS13 co-expressed with AurS12 (no His-tag). (A) Size exclusion chromatography (B) SDS-PAGE gel of Ni-NTA purification of His-tagged AurS13 expressed alone and His-tagged AurS13 co-expressed with AurS12 (no His-tag) (L: ladder, WCL: wash collection, E: Elution)

AurS12 and AurS13-$His_6$ were expressed and purified by Ni-NTA purification and anionic separation (FPLC, intact proteins), we observed a higher protein yield of AurS13 when co-expressed with AurS12. Size exclusion chromatography of AurS13 also showed a change from oligomers to dimer formation when the protein was co-expressed with AurS12. This is consistent with a previous study reporting conformation changes of DesVII in the presence of DesVIII (Borisova 2010).

**Table S2.** $^1$H and $^{13}$C NMR data of auroramycin analog **4** and analog **5**.

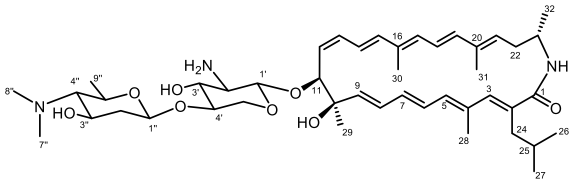

**4**

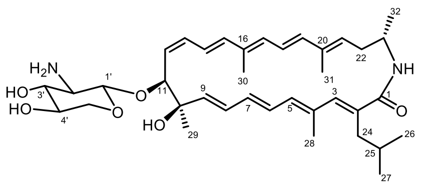

**5**

| | **4** ($CD_2Cl_2$:$CD_3OD$ = 5:1) | | **5** (DMSO-$d_6$) | |
|---|---|---|---|---|
| **No.** | **$\delta_C$** | **$\delta_H$ (multiplicity, *J* (Hz))** | **$\delta_C$** | **$\delta_H$ (multiplicity, *J* (Hz))** |
| 1 | 177.30, C | | 174.57,170.44,C | |
| 2 | 135.67, C* | | 135.41, C* | |
| 3 | | 137.42, CH | 136.19, CH | 6.14 (1H, s) |
| 4 | 133.56, C | | 131.97, C | |
| 5 | 136.15, CH | 5.78 (1H, m) | 134.65, CH | 5.70 (1H, d, 11.5) |
| 6 | 128.63, CH | 6.39 (1H, m) | 127.44, CH* | 6.45 (1H, m) |
| 7 | 135.67, CH* | 5.90 (1H, m) | 134.87, CH | 5.79 (1H, t, 12.8) |
| 8 | 130.47, CH | 6.21 (1H, m) | 130.87, CH | 6.12 (1H, m) |
| 9 | 140.95, CH | 5.51 (1H, m) | 142.23, CH | 5.47 (1H, d, 15.6) |
| 10 | 76.33, C | | 75.51, C | |
| 11 | 83.61, 88.89 CH | 4.25 (1H, m) | 82.29, CH | 4.24 (1H, d, 8.3) |
| 12 | 130.17, CH | 5.49 (1H, m) | 130.73, CH | 5.40 (1H, m) |
| 13 | 129.39, CH | 6.16 (1H, m) | 127.44, CH* | 6.16 (1H, m) |
| 14 | 125.57, CH | 6.19 (1H, m) | 125.56, CH | 6.15 (1H, m) |
| 15 | 138.11, CH | 6.28 (1H, m) | 135.41, CH* | 6.20 (1H, m) |
| 16 | 134.17, C | | 133.35, C | |
| 17 | 132.02, CH | 6.11 (1H, m) | 127.54, CH | 6.00 (1H, m) |
| 18 | 124.45, CH | 6.33 (1H, m) | 123.14, CH | 6.30 (1H, m) |
| 19 | 137.64, CH | 6.39 (1H, m) | 137.24, CH | 6.22 (1H, m) |
| 20 | 136.64, C | | 137.69, C | |
| 21 | 129.92, CH | 5.58 (1H, m) | 127.27, CH | 5.68 (1H, m) |
| 22 | 39.52, $CH_2$ | 1.12 (1H, m)<br>1.47 (1H, m) | 35.84, $CH_2$ | 2.42 (2H, m) |
| 23 | 46.88, 45.26 CH | 3.97 (1H, m) | 45.13, CH | 3.93 (1H, m) |
| 24 | 34.76, $CH_2$ | 2.23 (1H, t, 7.6)<br>2.49 (1H, m) | 33.75, $CH_2$ | 2.17 (1H, m) |
| 25 | 29.14, CH | 1.61 (1H, m) | 24.55, CH | 1.57 (1H, m) |
| 26 | 22.46, $CH_3$ | 0.83 (3H, m)^ | 22.29, $CH_3$ | 0.82 (3H, m) |
| 27 | 22.54, $CH_3$ | 0.83 (3H, m)^ | 22.34, $CH_3$ | 0.84 (3H, d, 6.7) |
| 28 | 16.38, $CH_3$ | 1.96 (3H, s) | 16.09, $CH_3$ | 1.94 (3H, s) |
| 29 | 20.96, $CH_3$ | 1.23 (3H, br s)^ | 23.62, $CH_3$ | 1.41 (3H, brs) |
| 30 | 12.99, $CH_3$ | 1.71 (3H, s) | 12.72, $CH_3$ | 1.71 (3H, s) |
| 31 | 12.87, $CH_3$ | 1.73 (3H, s) | 11.23 $CH_3$ | 0.80 (3H, m) |
| 32 | 19.31, $CH_3$ | 1.23 (3H, br s)^ | 20.81, $CH_3$ | 1.17 (3H, d, 6.7) |
| 1-NH | | | | |
| 1' | 104.60, 106.80, CH | 4.29 (1H, m) | 105.37, CH | 4.18 (1H, d, 7.6) |
| 2' | 57.67, CH | 2.68 (1H, m) | 57.74, CH | 2.49 (1H, m) |
| 3' | 74.99, CH | 3.31 (1H, m) | 76.00, CH | 3.02 (1H, m) |
| 4' | 79.69, CH | 3.53 (1H, m) | 69.59, CH | 3.26 (1H, m) |
| 5' | 64.32, $CH_2$ | 3.22 (1H, t, 11.0)<br>3.89 (1H,dd, 5.4, 11.8) | 66.00, $CH_2$ | 3.00 (1H, m)<br>3.66 (1H, dd, 5.2, 11.4) |
| 1" | 100.27, CH | 4.48 (1H, dd, 2.0, 9.8) | | |
| 2" | 39.79, $CH_2$ | 1.48 (1H, m)<br>2.20 (1H, ddd, 2.0, 4.8, 12.3) | | |
| 3" | 71.05, CH | 3,52 (1H, m) | | |
| 4" | 72.04, CH | 2.02 (1H, t, 9.8) | | |
| 5" | 66.24, CH | 3.62 (1H, ddd, 4.8, 9.8, 11.3) | | |
| 6" | | | | |
| 7" | 41.55, $CH_3$* | 2.40 (3H, s)^ | | |
| 8" | 41.55, $CH_3$* | 2.40 (3H, s)^ | | |
| 9" | 19.96, $CH_3$ | 1.28 (3H, d, 6.2) | | |

***overlap $^{13}$C signals; ^overlap $^1$H signals; chemical shifts in ppm using $CD_2Cl_2$ ($\delta_H$ = 5.32 ppm, $\delta_C$ = 53.84 ppm) or DMSO-6 ($\delta_H$ = 2.50 ppm, $\delta_C$ = 39.52 ppm) as reference.**

**Table S3.** $^{1}H$ and $^{13}C$ NMR data of auroramycin analog **10**.

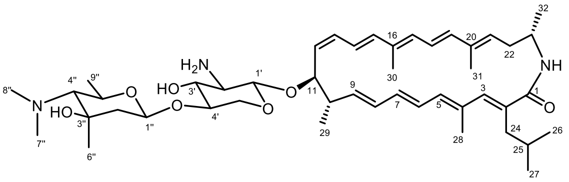


**10**

| No. | **10** ($CD_2Cl_2$:$CD_3OD$ = 5:1) δC | δH (multiplicity, *J* (Hz)) |
|---|---|---|
| 1 | 173.00, 177.15, C | |
| 2 | 136.52, C | |
| 3 | 136.89, CH | 6.17 (1H, s) |
| 4 | 132.40, C | |
| 5 | 136.78, CH | 5.80 (1H, m) |
| 6 | 126.82, CH | 6.33 (1H, m)^ |
| 7 | 136.52, CH | 5.91 (1H, m) |
| 8 | 131.69, CH | 6.16 (1H, m) |
| 9 | 140.04, CH | 5.61 (1H, m) |
| 10 | 28.41, CH | 1.48 (1H, m) |
| 11 | 70.66, CH | 3.89 (1H, m) |
| 12 | 130.06, CH | 5.44 (1H, m) |
| 13 | 129.18, CH | 6.14 (1H, m) |
| 14 | 124.57, CH | 6.42 (1H, m) |
| 15 | 137.30, CH | 6.33 (1H, m)^ |
| 16 | 133.55, C | |
| 17 | 131.44, CH | 7.02 (1H, m) |
| 18 | 115.65, CH | 6.73 (1H, m) |
| 19 | 137.46, CH | 6.40 (1H, m) |
| 20 | 137.40, C | |
| 21 | 136.68, CH | 6.20 (1H, m) |
| 22 | 30.10, $CH_2$ | 1.24 (2H, m)^ |
| 23 | 46.86, CH* | 3.96 (1H, m)^ |
| 24 | 36.94, $CH_2$ | 2.24 (1H, m) |
| | | 2.48 (1H, m) |
| 25 | 29.29, CH | 1.62 (1H, m) |
| 26 | 22.53, $CH_3$ | 0.84 (3H, m) |
| 27 | 22.73, $CH_3$ | 0.85 (3H, m) |
| 28 | 16.33, $CH_3$ | 1.96 (3H, s) |
| 29 | 23.10, $CH_3$ | 0.83 (3H, m) |
| 30 | 12.90, $CH_3$ | 1.73 (3H, s) |
| 31 | 12.97, $CH_3$ | 1.75 (3H, s) |
| 32 | 23.32, $CH_3$ | 1.24 (3H, m)^ |
| 1-NH | | |
| 1’ | 104.76, CH | 4.23 (1H, m) |
| 2’ | 57.51, CH | 2.71 (1H, m) |
| 3’ | 74.84, CH | 3.31 (1H, m) |
| 4’ | 79.84, CH | 3.52 (1H, m) |
| 5’ | 64.34, $CH_2$ | 3.20 (1H, t, 11.3) |
| | | 3.87 (1H, m) |
| 1” | 99.74, CH | 4.55 (1H, d, 9.9) |
| 2” | 45.83, $CH_2$ | 1.60 (1H, m) |
| | | 1.87 (1H, m) |
| 3” | 71.96, C | |
| 4” | 74.42, CH | 2.24 (1H, m) |
| 5” | 69.97, CH | 3.72 (1H, m) |
| 6” | 23.10, $CH_3$ | 1.24 (3H, s)^ |
| 7” | 44.04, $CH_3$* | 2.44 (3H, s)^ |
| 8” | 44.04, $CH_3$* | 2.44 (3H, s)^ |
| 9” | 20.91, $CH_3$ | 1.30 (3H, d, *J* = 5.9 Hz) |

***overlap $^{13}C$ signals; ^overlap $^{1}H$ signals; chemical shifts in ppm using $CD_2Cl_2$ ($\delta_H$ = 5.32 ppm, $\delta_C$ = 53.84 ppm) as reference.**

**Figure S11.** Analytical data for the structural assignment of auroramycin analog **3**.

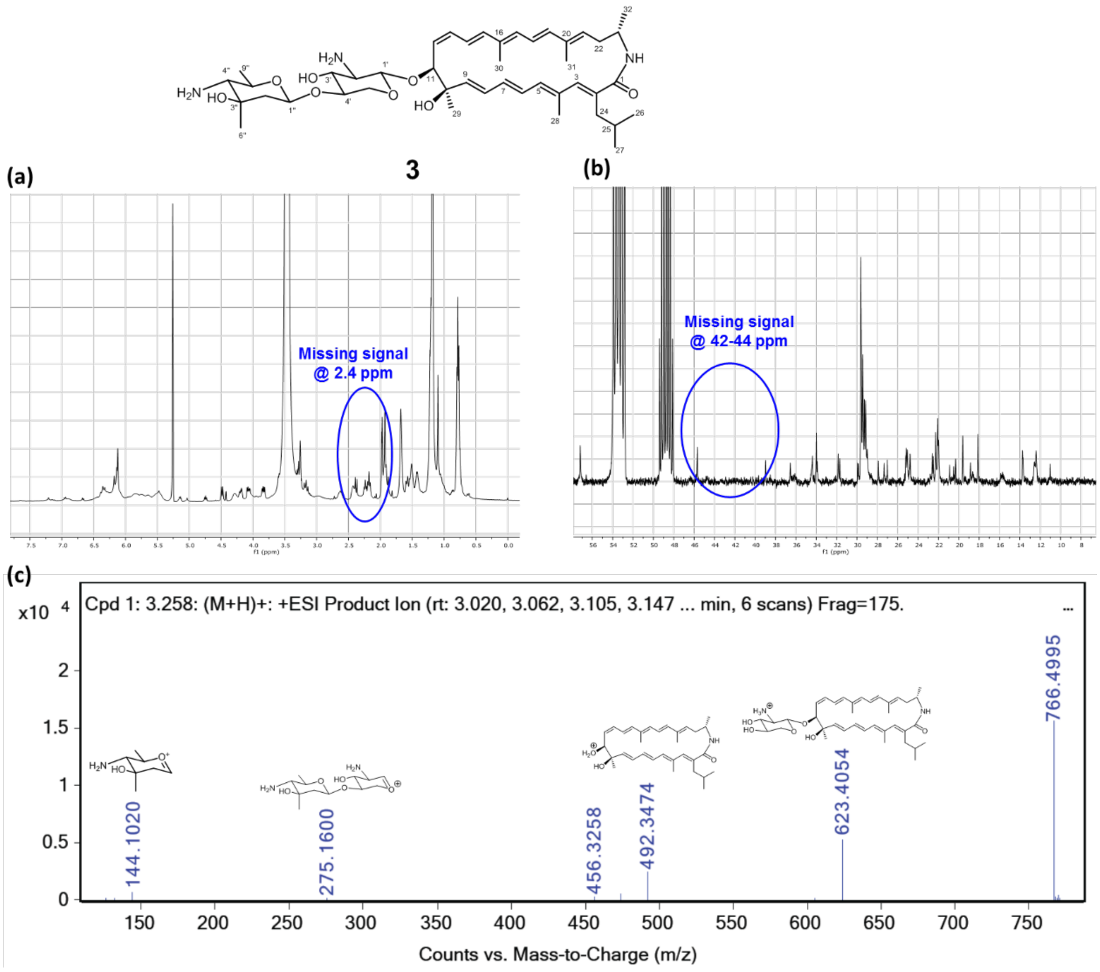


Structural assignment of auroramycin analog **3** was confirmed by combined inference of the NMR, HRMS and MS/MS data. Key features from the $^{1}$H and $^{13}$C that corroborate with the assigned structure are the disappearance of the methyl $\boldsymbol{\delta_H}$ and $\boldsymbol{\delta_c}$ signals at around 2.4 ppm and 42-44 ppm respectively (Figure S11a and S11b). This is further supported by the MS/MS data in which the MS2 signal 144.1020 can be assigned to the *N*,*N*-demethylated 3, 5-*epi*-lemonose unit (Figure S11c).

**Figure S12.** Analytical data for the structural assignment of auroramycin analog **6.**

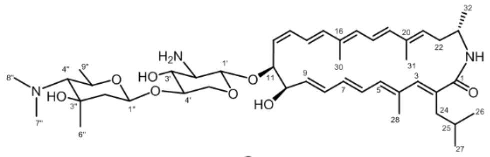

**6**

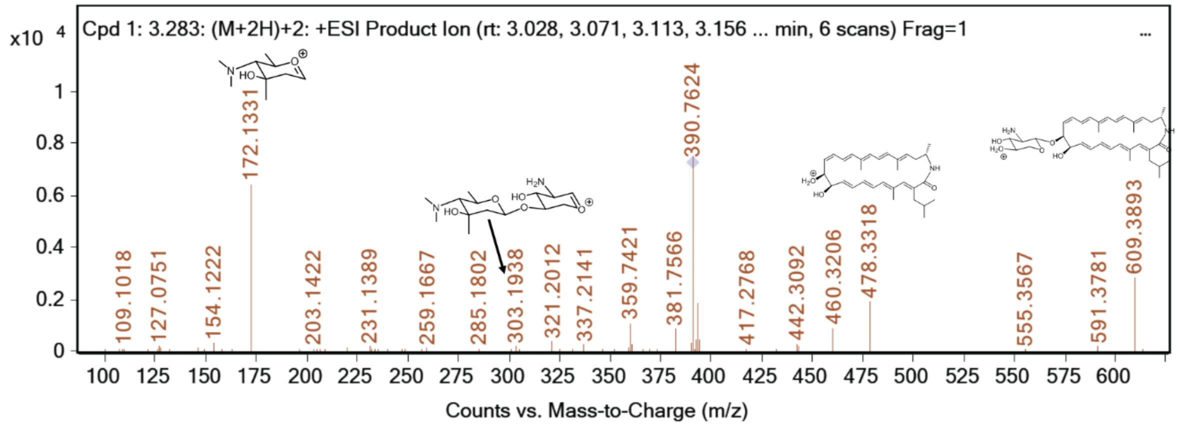


Structural assignment of auroramycin analog **6** was confirmed by inference of the HRMS and MS/MS data. Key features of the MS2 spectra has been assigned. MS2 signal 478.3318 can be assigned to the macrolactam fragment which suggests a methyl group is missing in analog **6** compared to auroramycin.

**Figure S13.** Analytical data for the structural assignment of auroramycin analog **7.**

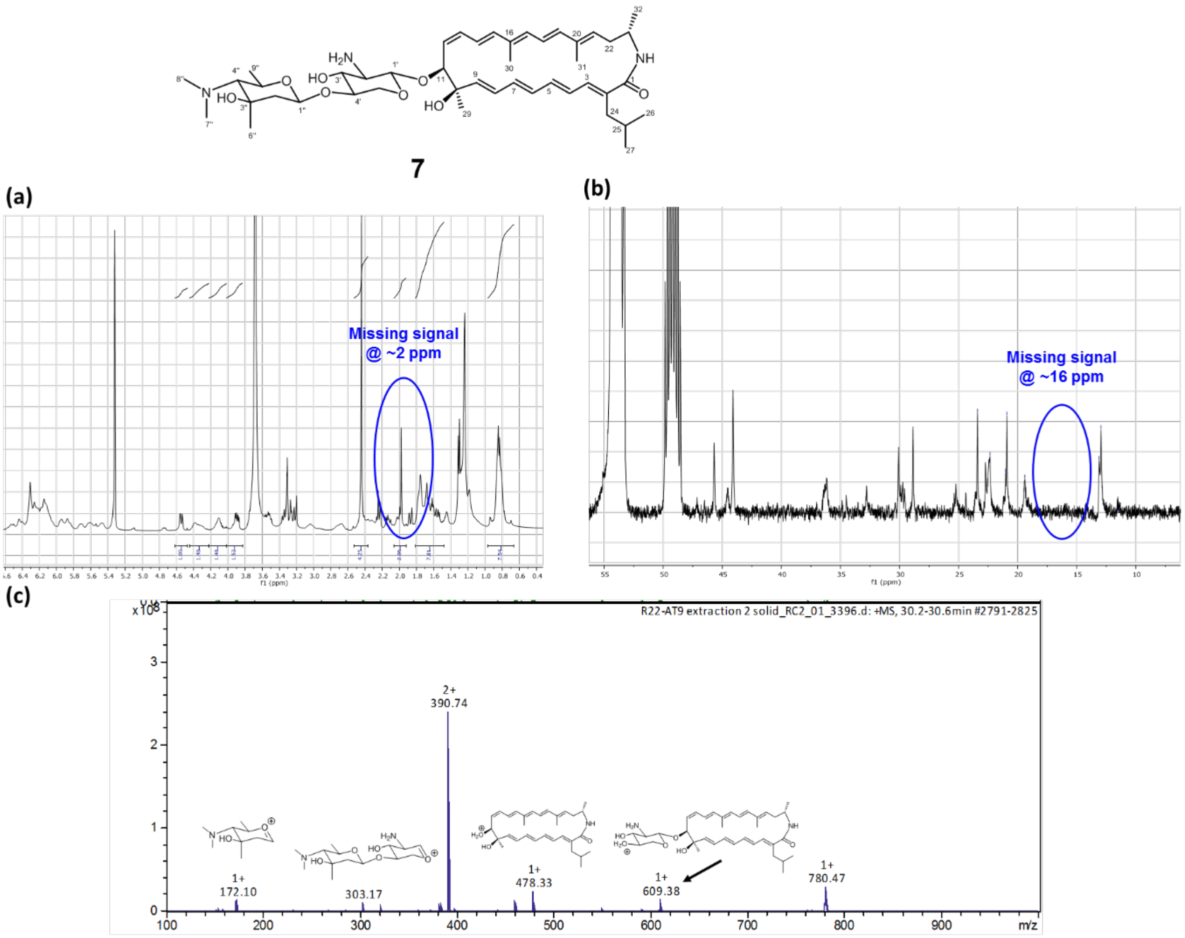


Structural assignment of auroramycin analog **7** was confirmed by combined inference of the NMR, HRMS and MS/MS data. Key features from the $^{1}H$ and $^{13}C$ that corroborate with the assigned structure are the disappearance of the methyl $\delta_H$ and $\delta_c$ signals at around 2 ppm and 16 ppm respectively (Figure S13a and S13b). This is further supported by the MS/MS data in which the MS2 signal 478.33 can be assigned to the macrolactam fragment which suggests a methyl group is missing in analog **7** compared to auroramycin (Figure S13c).

### References:

Moncrieffe MC, Fernandez MJ, Spiteller D, Matsumura H, Gay NJ, Luisi BF, Leadlay PF. Structure of the glycosyltransferase EryCIII in complex with its activating P450 homologue EryCII. Journal of molecular biology. 2012 Jan 6;415(1):92-101.

Borisova SA, Liu HW. Characterization of glycosyltransferase DesVII and its auxiliary partner protein DesVIII in the methymycin/pikromycin biosynthetic pathway. Biochemistry. 2010 Aug 24;49(37):8071-84.
